# Mpe1 senses the polyadenylation signal in pre-mRNA to control cleavage and polyadenylation

**DOI:** 10.1101/2021.09.02.458805

**Authors:** Juan B. Rodríguez-Molina, Francis J. O’Reilly, Eleanor Sheekey, Sarah Maslen, J. Mark Skehel, Juri Rappsilber, Lori A. Passmore

## Abstract

Most eukaryotic messenger RNAs (mRNAs) are processed at their 3’-end by the cleavage and polyadenylation factor (CPF/CPSF). CPF mediates endonucleolytic cleavage of the pre-mRNA and addition of a polyadenosine (poly(A)) tail, which together define the 3’-end of the mature transcript. Activation of CPF is highly regulated to maintain fidelity of RNA processing. Here, using cryoEM of yeast CPF, we show that the Mpe1 subunit directly contacts the polyadenylation signal sequence in nascent pre-mRNA. This RNA-mediated link between the nuclease and polymerase modules promotes activation of the CPF endonuclease and controls polyadenylation. Mpe1 rearrangement is antagonized by another subunit, Cft2. *In vivo*, depletion of Mpe1 leads to widespread defects in transcription termination by RNA Polymerase II, resulting in transcription interference on neighboring genes. Together, our data suggest that Mpe1 plays a major role in selecting the cleavage site, activating CPF and ensuring timely transcription termination.

## Introduction

Co-transcriptional processing of pre-mRNAs is crucial for their nuclear export, cellular localization, stability and translation (Moore and Proudfoot, 2009). Processing is carried out by a series of multi-subunit enzymatic complexes that are recruited to RNA polymerase II (RNAPII) throughout the transcription cycle to mediate 5’-capping, splicing, and 3’-end processing (Hocine et al., 2010). Crosstalk between these events safeguards transcription homeostasis (Moore and Proudfoot, 2009).

mRNA 3’-end processing involves a specific endonucleolytic cleavage of the pre-mRNA, and addition of a polyadenosine (poly(A)) tail onto the new 3’-end by the cleavage and polyadenylation factor (CPF in yeast, CPSF in human) (Kumar et al., 2019; Sun et al., 2020). Endonucleolytic cleavage releases pre-mRNA from the site of transcription and creates an exposed 5’ monophosphate in the RNAPII-bound nascent RNA. The unprotected 5’-end serves as a substrate for the torpedo exonuclease (Rat1), which degrades the downstream RNA and displaces RNAPII from chromatin, promoting transcription termination (Kim et al., 2004). Thus, regulated cleavage by CPF/CPSF not only defines the 3ʹ-UTR sequence of the mRNA, it is also required for transcription termination. Premature activation of pre-mRNA cleavage results in aberrant transcripts, whereas delayed activation causes transcriptional readthrough.

CPF subunits, and 3’-end processing in general, are highly conserved across all eukaryotes. In the yeast *Saccharomyces cerevisiae*, CPF is assembled into a 14 subunit (∼850 kDa) complex, that is organized into three enzymatically distinct and interconnected modules; the polymerase, nuclease and phosphatase modules (Casanal et al., 2017).

The polymerase module (mammalian polyadenylation specificity factor, or mPSF, in human) serves as the central interaction hub for the 3’-end processing machinery and harbors the poly(A) polymerase, Pap1. CryoEM structures of the core polymerase module from yeast and human revealed an assembly of four beta propellers within Cft1 and Pfs2 (CPSF160 and WDR33 in human) (Casanal et al., 2017; Clerici et al., 2017; Clerici et al., 2018; Sun et al., 2018). These act as a scaffold for the RNA-binding subunit Yth1 and the Pap1-binding subunit Fip1 (CPSF30 and FIP1 in human) (Casanal et al., 2017).

The nuclease module of CPF contains the endonuclease Ysh1, the pseudo- nuclease Cft2, and the multidomain protein Mpe1 (orthologs of human CPSF73, CPSF100 and RBBP6, respectively). The nuclease module as a whole appears to be flexible with respect to the polymerase module (Hill et al., 2019; Zhang et al., 2020). In human, the two modules are tethered together via a conserved peptide motif (mPSF interaction motif, or PIM) in CPSF100 that interacts with a surface groove on CPSF160 (Zhang et al., 2020).

Mpe1 is a stable component of yeast CPF (Vo et al., 2001). A conserved N-terminal ubiquitin-like domain (UBL) in Mpe1 interacts with the nuclease domain of Ysh1 (Hill et al., 2019). Additional domains in Mpe1, including a zinc knuckle and RING finger, are thought to interact with RNA (Lee and Moore, 2014). *In vivo* crosslinking experiments suggest that Mpe1 contacts nascent pre-mRNAs very close to the cleavage site (∼6 nucleotides upstream) with a preference for A/U rich sequences (Baejen et al., 2014). Functionally, Mpe1 and RBBP6 stimulate both cleavage and polyadenylation, and help select the cleavage site (Di Giammartino et al., 2014; Lee and Moore, 2014; Vo et al., 2001). RBBP6 co-immunoprecipitates with human CPSF (Di Giammartino et al., 2014) but their association may be transient. The mechanistic details regarding the precise role of Mpe1 and RBBP6 in 3ʹ-end processing remain unknown.

The phosphatase module includes two protein phosphatases, Ssu72 and Glc7 (orthologs of human SSU72 and PP1). Both of these phosphatases regulate transcription by dephosphorylating the C-terminal domain of the Rpb1 subunit of RNAPII (Jeronimo et al., 2016), for example to promote transcription termination (Schreieck et al., 2014).

Activation of the CPF endonuclease is highly controlled and requires coordinated assembly of CPF with two accessory RNA-binding factors, cleavage factors IA and IB (CF IA and CF IB) (Gordon et al., 2011; Gross and Moore, 2001; Hill et al., 2019; Kessler et al., 1997; Kumar et al., 2019). CPF, CF IA and CF IB each bind to specific sequence elements in pre-mRNAs and together they activate the 3ʹ-end processing machinery.

The polyadenylation signal (PAS), also referred to as the positioning element in yeast, is conserved from yeast to mammals with a consensus sequence of A1A2U3A4A5A6 (Guo and Sherman, 1996; Russo et al., 1993; Tian and Graber, 2012). In mammals, the mechanistic basis of PAS recognition is well understood. Structures of mPSF show that CPSF30 zinc finger 2 binds A1 and A2 of the PAS, and zinc finger 3 binds A4 and A5 (Clerici et al., 2018; Sun et al., 2018). U3 and A6 form a Hoogsteen base-pair which inserts into a hydrophobic pocket of WDR33 (Clerici et al., 2018; Sun et al., 2018). In yeast the PAS sequence is more degenerate but Yth1 zinc fingers 2 and 3 are predicted to interact with the A1A2 and A4A5 di-nucleotides of the yeast PAS similar to human. In contrast, the N-terminal loop of WDR33 that binds the U3:A6 Hoogsteen base pair is not conserved in the yeast counterpart, Pfs2. Finally, how PAS recognition results in endonuclease activation in yeast and human remains unanswered.

Here, we present structural, biochemical and transcriptomic evidence that Mpe1 binds the polymerase module and that, surprisingly, Mpe1 makes direct contact with the PAS RNA. Mpe1 interaction with both the polymerase module-RNA complex and Ysh1 activates cleavage, and regulates polyadenylation by CPF. Acute depletion of Mpe1 in yeast causes widespread defects in transcription termination, which leads to transcription interference. Overall, our work suggests that Mpe1 senses RNA, directly linking RNA binding by CPF with regulation of cleavage, polyadenylation and transcription termination.

## Results

### Mpe1 interacts directly with the polymerase module

To gain insight into the function of Mpe1, we first investigated whether it links the nuclease and polymerase modules through a direct interaction with any of the five subunits of the polymerase module. After pairwise co-expression in insect cells, Cft1 (but no other polymerase module subunits) co-purified with StrepII-tagged Mpe1 (Mpe1- SII) (Figure S1A). This is consistent with previous data that suggested Mpe1 interacts with Cft1 (Lee and Moore, 2014). We also tested for Mpe1 interaction with purified polymerase module and found that they formed a complex that could be purified by size exclusion chromatography, although Mpe1 was associated at substoichiometric levels (Figure S1B).

An Mpe1 construct including the zinc knuckle and the downstream linker was previously shown to interact with Cft1 in a yeast-two-hybrid assay (Lee and Moore, 2014). To identify regions in Mpe1 that may contact the polymerase module in a fully reconstituted system, we performed hydrogen-deuterium exchange followed by mass spectrometry (HDX-MS). This method identified Mpe1 peptides that are protected from solvent exchange upon interaction with the polymerase module. HDX-MS showed protection within several regions that include the zinc knuckle, the RING finger and a C-terminal region (Figure S1C), indicating that Mpe1 may make multiple contacts with the polymerase module, or that Mpe1 may undergo a major rearrangement upon binding.

Because Mpe1 had previously been implicated in RNA binding (Baejen et al., 2014; Lee and Moore, 2014), we next tested whether the polymerase module and Mpe1 form a stable complex with RNA. For these experiments, we used the 3ʹ-end of the *CYC1* transcript that is often used as a model substrate for *in vitro* cleavage and polyadenylation assays (Hill et al., 2019). Specifically, we used a 5ʹ FAM-labelled synthetic RNA that corresponds to the 5ʹ product of the cleavage reaction of the *CYC1* 3ʹ UTR, and includes the AAGAA PAS sequence (see Figure 2A below). We refer to this 42 nt RNA as pre-cleaved *CYC1* RNA. In size exclusion chromatography, the polymerase module, Mpe1 and RNA co-migrated, suggesting that they form a complex (Figure S1B). RNA promoted a more stoichiometric association of Mpe1 with the polymerase module, suggesting that RNA stabilizes Mpe1 on the complex. Interestingly, RNA shifted the polymerase module-Mpe1 complex to a later elution volume, consistent with potential compaction of the complex. Free Mpe1 also co-eluted with RNA confirming a direct association of Mpe1 with RNA. Together, these data suggest that in the presence of RNA, Mpe1 interaction with the polymerase module is stabilized, and the complex may undergo a conformational change that results in its compaction.

Next, we analyzed the polymerase module-Mpe1-RNA complex by single particle electron cryo-microscopy (cryoEM) (Table 1, Figure S1D-E). Compared to the previous structure of yeast polymerase module (Casanal et al., 2017), the sample we used here additionally contained Pap1, Mpe1 and the pre-cleaved *CYC1* RNA. We were able to obtain a map of the complex at an overall resolution of 2.67 Å (Figures 1A and S1F-I).

**Figure 1.**
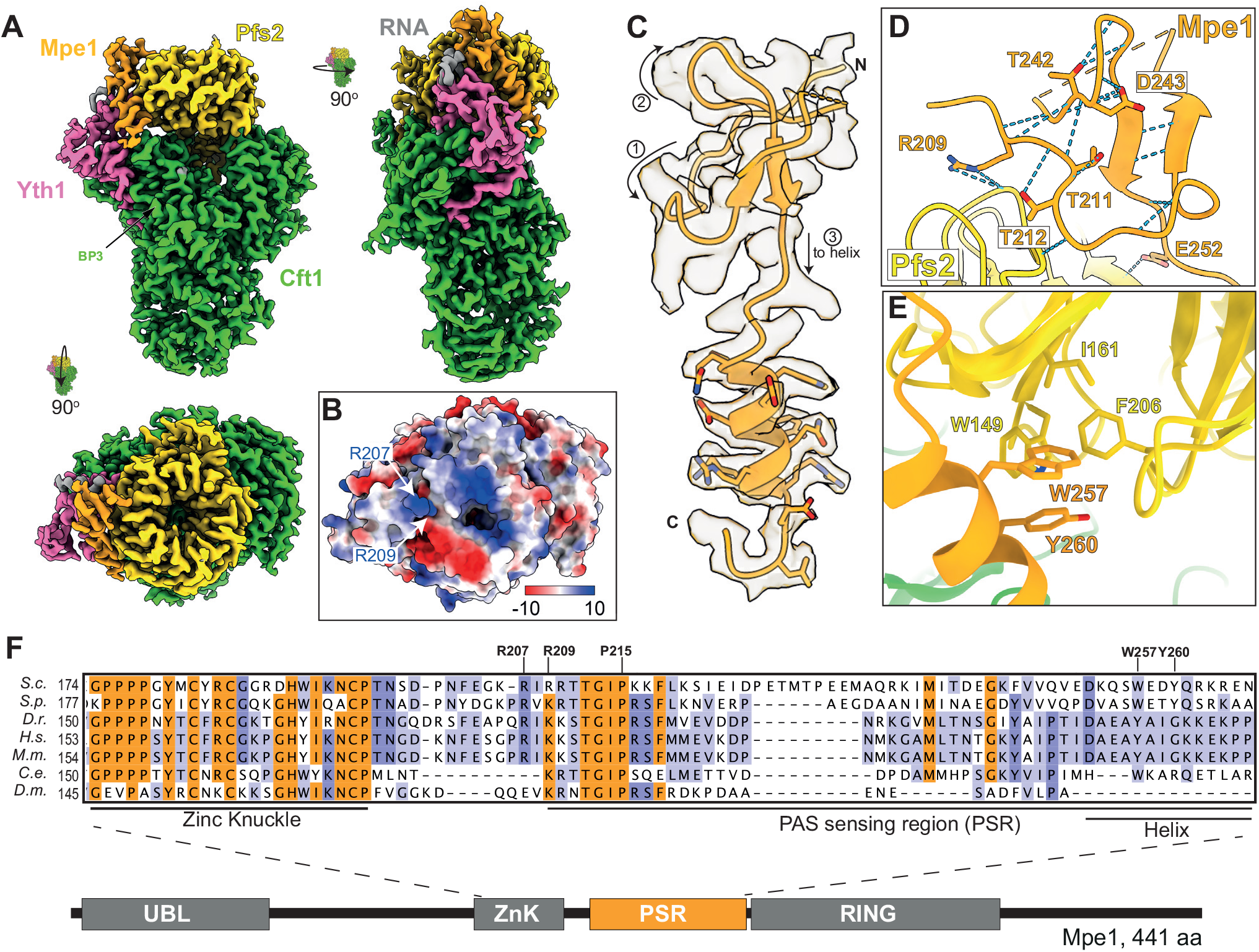
Structure of Mpe1 bound to the polymerase module of CPF. (A) Front, side and top views of the cryoEM map of the polymerase module (Cft1; green, Pfs2; yellow, Yth1; pink) in complex with Mpe1 (orange) and RNA (grey). Beta-propeller 3 (BP3) of Cft1 is indicated. (B) Surface representation of top view of the model of polymerase module-Mpe1-RNA (looking down the center of the beta-propeller of Pfs2), colored by electrostatic potential (±10 kT/e). Highlighted residues (R207 and R209) belong to Mpe1. (C) Cartoon representation of residues 207–268 of the Mpe1 polyadenylation signal-sensing region (PSR) within its corresponding section of the cryoEM map. The direction of the polypeptide chain is shown with arrows and numbered 1 to 3. The N- and C-termini are labeled. (D) Hydrogen bond network (blue dashed lines) within Mpe1 residues 207–252. Side chains involved in hydrogen bonds are shown in sticks; all other hydrogen bonds are with main chain atoms. In (C) and (D), orange dashes denote a disordered region of the map (residues 224–239). (E) Selected residues of the Mpe1 PSR helix (orange, W257 and Y260) and the hydrophobic pocket of Pfs2 (yellow). (F) Multiple sequence alignment of the zinc knuckle and PSR of Mpe1 orthologues. Residues highlighted in orange are conserved across most eukaryotes; those in purple are partially conserved. A domain diagram of Mpe1 is shown below. *S.c; Saccharomyces cerevisiae*, *S.p. Schizosaccharomyces pombe*, *D.r. Danio rerio*, *H.s. Homo sapiens*, *M.m. Mus musculus*, *C.e. Caenorhabditis elegans*, *D.m. Drosophila melanogaster*. See also Figures S1 and S2.

**Table 1.** CryoEM data collection, model refinement and validation statistics.

|  | polymerase<br>module-Mpe1-RNA | polymerase<br>module-Cft2(S) | polymerase<br>module-Mpe1-yPIM-RNA |
| --- | --- | --- | --- |
| **Data collection and processing** |  |  |  |
| Magnification | 105,000 x | 105,000 x | 105,000 x |
| Voltage (keV) | 300 | 300 | 300 |
| Electron exposure (e<sup>-</sup>/Å<sup>2</sup>) | 40 | 37 | 40 |
| Defocus range (um) | -0.5 to 3.1 | -0.5 to 3.1 | -0.5 to 3.1 |
| Pixel size (Å) | 0.83 (eBIC) | 0.86 (LMB) | 0.86 (LMB) |
| Symmetry imposed | C1 | C1 | C1 |
| Initial particle images (no.) | 6,460,073 | 1,946,027 | 13,905,256 |
| Final particle images (no.) | 131,152 | 141,584 | 846,349 |
| Map resolution (Å) | 2.66 | 2.79 | 2.61 |
| FSC threshold | 0.143 | 0.143 | 0.143 |
| Map resolution range (Å) | 2.66 to >10 | 2.79 to >10 | 2.61 to >10 |
| **Refinement** |  |  |  |
| Initial model used | *De novo*<br>modelling and<br>polymerase<br>module (6eoj) | mPSF-PIM<br>(6urg) and<br>polymerase<br>module (6eoj) | Polymerase<br>module-Mpe1-RNA and<br>polymerase<br>module-yPIM |
| Model resolution (Å) |  |  |  |
| FSC threshold | 0.143 | 0.143 | 0.143 |
| Model resolution range (Å) | n/a | n/a | n/a |
| Map sharpening *B* factor (Å<sup>2</sup>) | -20 | -30 | -40 |
| **Model composition** |  |  |  |
| Non-hydrogen atoms | 14,207 | 14,027 | 14,616 |
| Protein residues | 1,768 | 1,754 | 1,819 |
| Nucleotides | 3 | 0 | 3 |
| Ligands | ZN:2 | ZN:2 | ZN:2 |
| *B* factors (Å<sup>2</sup>) |  |  |  |
| Protein | not estimated | not estimated | not estimated |
| Ligand |  |  |  |
| **r.m.s. deviations** |  |  |  |
| Bond lengths (Å) | 0.003 | 0.008 | 0.003 |
| Bond angles (°) | 0.532 | 1.258 | 0.537 |
| **Validation** |  |  |  |
| MolProbity score | 1.93 | 1.72 | 1.85 |
| Clashscore | 9.25 | 4.50 | 7.93 |
| Poor rotamers (%) | 0 | 0 | 0.06 |
| **Ramachandran plot** |  |  |  |
| Favoured (%) | 93.12 | 91.73 | 93.67 |
| Allowed (%) | 6.77 | 8.09 | 6.16 |
| Disallowed (%) | 0.11 | 0.17 | 0.17 |

In this map, we identified an additional density, not present in previous maps of the yeast polymerase module or mPSF (Casanal et al., 2017; Zhang et al., 2020), that extends from the top of Pfs2 towards Yth1 and Cft1 (Figure 1A, orange). There is also poorly ordered density on top of the Pfs2 beta propeller and in front of the C-terminal helical domain of Cft1 (Figure S2A).

Given the resolution of the map, we could *de novo* build an atomic model into part of the additional density, including a region of Mpe1 downstream of the zinc knuckle (residues 209–222 and 240–268). This region of Mpe1 primarily contacts Pfs2 but also comes into close proximity of Cft1 and Yth1. Residues 223–239 are not ordered and could not be resolved in our structure. We identified the remaining well-ordered density as three nucleotides of the PAS of the *CYC1* RNA. We therefore refer to the region of Mpe1 that is ordered in our maps as the PAS sensing region (PSR).

Densities for Fip1 and Pap1 were not identified in the map.

### Mpe1 interactions with Pfs2 and Cft1

The N-terminal part of the Mpe1 PSR is located near the top of Pfs2 (Figure 1A).

Two arginine residues in Mpe1 (R207 and R209) are positioned next to a positively-charged patch on the top surface of Pfs2 (Figure 1B), which was previously predicted to participate in RNA binding (Casanal et al., 2017). Thus, Mpe1 may also contribute to this putative RNA binding site. Interestingly, additional density over the positively-charged patch suggests that protein or RNA bind here, but this density is poorly ordered and we were not able to confirm its identity (Figure S2A).

Mpe1 residues 209–252 form a small, compact fold that packs against the Pfs2 beta-propeller. This fold is held together by a network of hydrogen bonds and a hydrophobic core (Figure 1C-D). Mpe1 then continues as a helix (residues 253–268) that makes additional contacts with the side of the Pfs2 beta-propeller. Two aromatic residues from this helix (W257 and Y260) insert into a hydrophobic pocket of Pfs2 that is lined by W149, I161, and F206 (Figure 1E). Mpe1 Y260 also forms a π-π interaction with W149 of Pfs2.

Mpe1 is not visible after residue Q268, which is positioned near beta-propeller 3 of Cft1. Overall, the Mpe1 PSR buries a combined surface area of ∼1,200 Å^2^ on the polymerase module, a relatively modest area that is consistent with the small number of residues (Chen et al., 2013) that stabilize the interaction between the PSR and the polymerase module. Curiously, although Mpe1 and Cft1 interact directly in pulldowns, there is little direct contact between them in the models. The additional density adjacent to the C-terminal alpha helical domain of Cft1 (Figure S2A) may represent an additional, flexible interaction between Mpe1 and Cft1. Together, these data reveal an unexpected architecture by which Mpe1 interacts with the polymerase module of CPF.

We aligned the sequences of Mpe1 orthologues from diverse eukaryotic species and found that PSR residues 209–218 are highly conserved (Figure 1F). For example, Mpe1 R209 and R210 near the positively-charged patch on Pfs2 are conserved as lysine or arginine. The loop that is not resolved in our map (residues 223–239) and the helix are not well conserved except for W257 which is conserved as an aromatic residue.

We also compared our model of yeast polymerase module-Mpe1-RNA to the structure of human mPSF. This showed that the Mpe1 PSR helix overlaps with a loop of CPSF30 (residues 22–34). Specifically, a phenylalanine (F30) of CPSF30 inserts into the hydrophobic pocket of WDR33 that binds Mpe1 W257 in yeast (Figure S2B). F30 of CPSF30 is conserved only among metazoans but the residues that line the hydrophobic pocket in Pfs2 are mostly conserved in WDR33 (W149/W175, I161/Y187 and F206/F231 in yeast/humans respectively) (Figure S2C-D) (Clerici et al., 2018). Thus, although some aspects of Mpe1 interaction with Pfs2 are conserved, the Mpe1 binding pocket is instead occupied by CPSF30 in human.

### Mpe1 senses the polyadenylation signal and stimulates polyadenylation

Three ribonucleotides fit into the density near zinc finger 2 of Yth1 (Figure 2A-B). These correspond to nucleotides A1 and A2 of the PAS of *CYC1* (A1A2G3A4A5), as well as one additional upstream nucleotide (U-1). There is weak density for a fourth nucleotide (G3). The first two nucleotides in the structure are arranged in a U-1-Yth1 Y83-A1 stack (Figure 2C). One face of the A2 base is stabilized by a π-π interaction with H69 of Yth1. This overall mechanism of binding to the A-dinucleotide by yeast Yth1 is very similar to its human counterpart, suggesting that recognition of the 5ʹ-end of the polyadenylation signal by zinc finger 2 is conserved (Figure S2E). In both human and yeast, sequence recognition is mediated by hydrogen bonds to the N1 amino groups of A1 and A2, and to the N6 amino group of A1. Consistent with the importance of RNA binding, zinc finger 2 is the most highly conserved Yth1 zinc finger across eukaryotes (Clerici et al., 2018).

**Figure 2.**
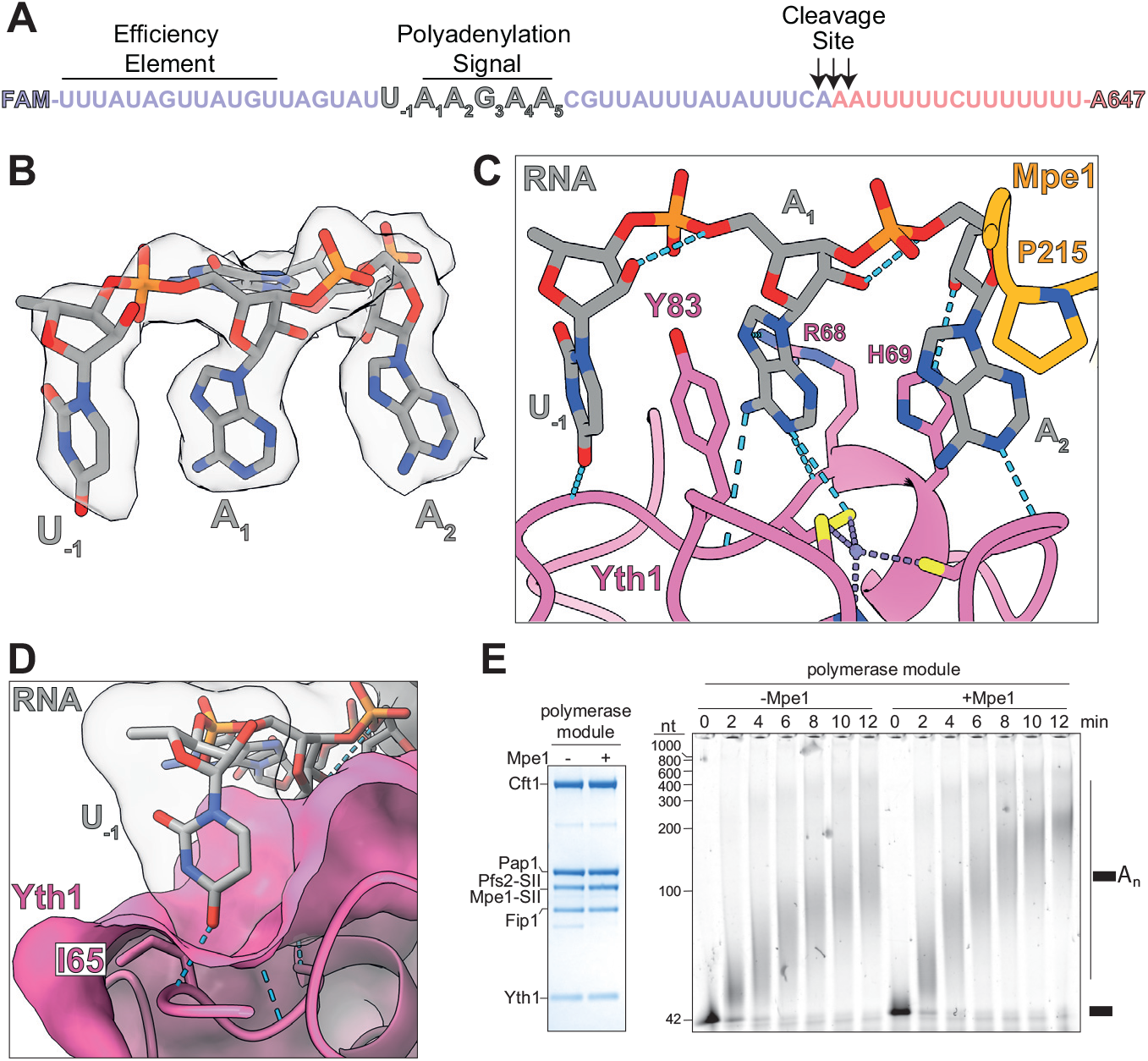
Mpe1 contacts the polyadenylation signal (PAS) in RNA and stimulates polyadenylation. (A) Sequence of the *CYC1* RNA substrate. Indicated are the 5ʹ cleavage product (purple, equivalent to the pre-cleaved *CYC1* substrate used in cryoEM and polyadenylation assays), the 3ʹ cleavage product (pink), the canonical cleavage sites (arrows), and the 5ʹ- and 3ʹ-fluorophores. The polyadenylation signal (PAS, also known as the positioning element) is indicated. (B) CryoEM map (transparent surface) and model (sticks) of RNA visible in the polymerase module-Mpe1-RNA map. Position numbering is relative to the sequence of the *CYC1* PAS. (C) Contacts between the PAS of *CYC1* (grey) with Yth1 (pink) and Mpe1 (orange). U-1 and A1 sandwich Y83 of Yth1, and R68 contacts the opposite face of the A1 base through a cation-π interaction. H69 of Yth1 stacks against one face of the A2, while P215 of Mpe1 contacts the opposite face via a CH-π interaction. Blue dashed lines show hydrogen bonds. (D) The U-1 nucleotide fits into an open pocket on Yth1 and hydrogen-bonds to the mainchain of I65. U-1 is shown in sticks representation inside its surface representation to illustrate the space it occupies in the pocket. (E) The effect of Mpe1 on polyadenylation activity of the polymerase module. Purified polymerase module (10 pmol) without or with Mpe1 was analyzed by SDS-PAGE and is shown on the left. Polyadenylation reactions using a 5ʹ FAM-labeled pre-cleaved *CYC1* RNA substrate were analyzed by urea-PAGE, shown on the right. CF IA and CF IB were not included in these reactions. The pre-cleaved *CYC1* RNA substrate is indicated with a black rectangle. See also Figure S2.

Surprisingly, in addition to the interactions between RNA and Yth1, A2 is contacted directly by P215 of Mpe1 through a CH-π interaction (Levitt and Perutz, 1988). P215 and the surrounding residues are conserved in Mpe1 homologs (Figure 1F), suggesting that this interaction may also be conserved across eukaryotes. Mpe1 is not required for polymerase module to form a complex with RNA (Figure S2F). We therefore hypothesize that Mpe1 senses RNA binding by Yth1.

The -1 nucleotide is not visible in previous mPSF-RNA structures. The base of U- 1 sits in an open pocket on the surface of Yth1, and there is a hydrogen-bond between the U-1 O4 and the main chain amide of I65 (Figure 2D). Interestingly, a *CYC1* RNA with a G in the -1 position is a poor cleavage substrate (Russo et al., 1993). The structure therefore suggests that the identity of the upstream nucleotide plays a role in RNA recognition by Yth1.

Given that Mpe1 contacts RNA and binds the polymerase module, we tested its effect on the polyadenylation activity of the polymerase module. We carried out *in vitro* polyadenylation assays using purified polymerase module with or without Mpe1, and the 5ʹ FAM-labeled pre-cleaved *CYC1* RNA as substrate. We then resolved the polyadenylated species on a denaturing urea gel. These data showed that Mpe1 promotes a modest but reproducible increase in the rate of polyadenylation (Figure 2E). The accessory factors CF IA and CF IB interact with the polymerase module and, in addition to activating cleavage, also increase the activity and processivity of polyadenylation (Casanal et al., 2017). In our assays, CF IA and CF IB mask the stimulatory effect of Mpe1 on polyadenylation (Figure S2G). Therefore, Mpe1 and cleavage factors may stimulate polyadenylation activity similarly by providing additional RNA binding sites on the polymerase module.

Overall, our structure reveals that the molecular mechanism of recognition of the first two nucleotides of the PAS by zinc finger 2 of Yth1 is highly-conserved across eukaryotes, and surprisingly, the PAS is also contacted by Mpe1. Our data also suggest that Mpe1 plays a role in stimulating the polyadenylation activity of the polymerase module.

### Cft2 antagonizes Mpe1 binding to polymerase module

To further understand how the nuclease and polymerase modules assemble, we added Cft2 to the polymerase module-Mpe1 complex. In size exclusion chromatography, Cft2 co-migrated with the polymerase module but, surprisingly, this reduced Mpe1 binding (Figures 3A and S3A, blue). Mpe1 incorporation was recovered by including the pre-cleaved *CYC1* RNA (Figures 3A and S3A, green). This suggests that there may be communication between Cft2, Mpe1 and RNA on the polymerase module.

**Figure 3.**
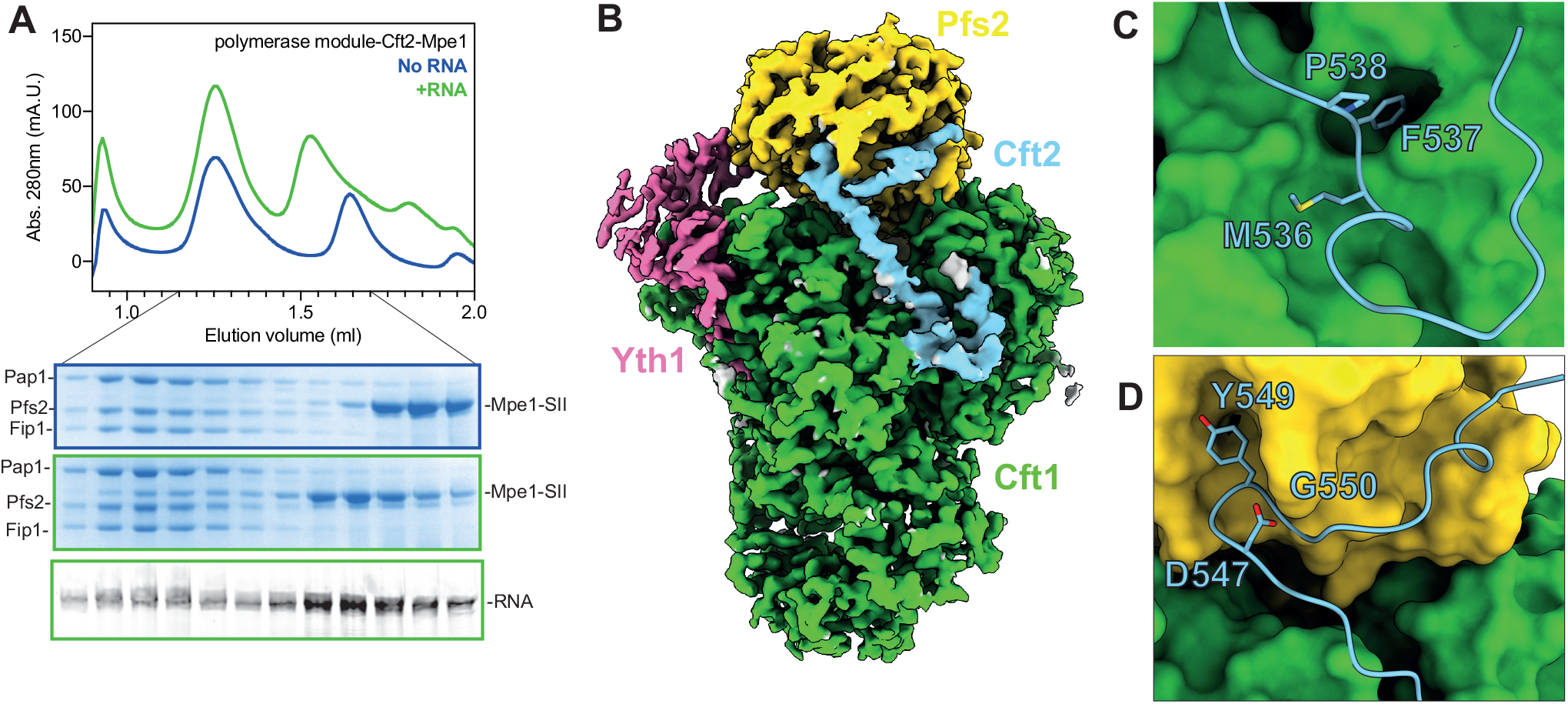
Cft2 antagonizes Mpe1 binding to polymerase module. (A) Size exclusion chromatography was performed with polymerase module in complex with Cft2 and Mpe1, with (green) or without (blue) pre-cleaved *CYC1* RNA. The chromatogram is shown in the top panel; Coomassie-stained SDS-PAGE of the indicated fractions are shown in the middle panels; and urea-PAGE of fluorescently-labeled RNA from the indicated fractions is shown in the bottom panel. The gels are outlined in colors corresponding to the chromatogram profiles. (B) CryoEM map of the polymerase module in complex with Cft2(S). The yeast polymerase module interacting motif (yPIM) of Cft2 is colored in blue. The rest of Cft2, Mpe1, pre-cleaved *CYC1* RNA, Fip1 and Pap1 were included in the sample but are not visible in the map. (C-D) The yPIM of Cft2 (blue, cartoon and sticks representation) inserts a highly-conserved F537 residue into a hydrophobic pocket in Cft1 (green, surface representation) (C), and a conserved Y549 residue into a hydrophobic pocket of Pfs2 (yellow, surface representation) (D). See also Figure S3.

The Mpe1 UBL domain interacts directly with the endonuclease subunit Ysh1, which in turn binds the C-terminal domain of Cft2 (Dominski et al., 2005; Hill et al., 2019). Thus, Ysh1 stabilizes Mpe1 association with the nuclease module via a mechanism that does not involve RNA. In agreement with this, Mpe1 co-eluted with the polymerase module in size exclusion chromatography in the presence of nuclease module subunits Ysh1 and Cft2, even in the absence of RNA (Figure S3B). We previously reported purification of this complex containing all subunits of the nuclease and polymerase modules, which we termed ‘CPFcore’ (Hill et al., 2019).

To determine the architecture of the polymerase module-Cft2-Mpe1-RNA complex, we carried out single particle cryoEM. We used a truncated version of Cft2 (Cft2(S), residues 1–720) which is missing the disordered C-terminal region but still interacts with CPF (Kyburz et al., 2003). This allowed us to purify a complex that was amenable to cryoEM sample preparation (Figure S3C). We obtained a map of this complex at an overall resolution of 2.79 Å (Figures 3B and S3D-G). This map contained density for the polymerase module and a short region of Cft2, but no density that corresponded to Mpe1, the pre-cleaved *CYC1* RNA, Fip1 or Pap1, despite their presence in the cryoEM specimen.

We built an atomic model of a conserved region of Cft2 that interacts with the polymerase module and is homologous to the human CPSF100 PIM (Zhang et al., 2020). We thus refer to this region of Cft2 (residues 525–562) as the yeast polymerase module interacting motif (yPIM). The yPIM adopts an arrangement on the polymerase module that is highly similar to its human counterpart on the mPSF (Figure S3H-I).

Conserved aromatic residues in the yPIM (F537 and Y549) stabilize its interaction with the polymerase module by inserting into hydrophobic pockets in Cft1 and Pfs2, respectively (Figure 3C-D). The yPIM also appeared to stabilize a loop in Cft1 (residues 1294–1303) that is normally disordered in its absence (Figure S3J).

The binding sites of the yPIM of Cft2 and the PSR of Mpe1 are non-overlapping.

This raised the question of how Cft2 antagonizes Mpe1 binding to the polymerase module (Figure 3A). To assess whether other parts of Cft2(S) interact with polymerase module or Mpe1, we performed photo-crosslinking using sulfo-NHS-diazirine (sulfo-SDA) followed by mass spectrometry analysis (Supplementary Information). We did not observe any crosslinks between Cft2 and Mpe1. However, a cluster of crosslinks confirm that the yPIM binds Cft1 in the groove between beta-propellers 1 and 3 near the Cft1 helical bundle (Figure 4A and B). Interestingly, we also detected crosslinks between Cft2 and Pfs2 (Figure 4A and B, yellow): Regions of Cft2 both upstream and downstream of the yPIM crosslink in the vicinity of the Mpe1 PSR binding site on Pfs2. Since we did not observe any Cft2 density in this region of the cryoEM map, Cft2 may occupy the space next to this part of Pfs2 without forming specific contacts. We conclude that full-length Cft2 is in close proximity to the Mpe1 binding site, suggesting that it may sterically clash with Mpe1 PSR binding to the polymerase module.

**Figure 4.**
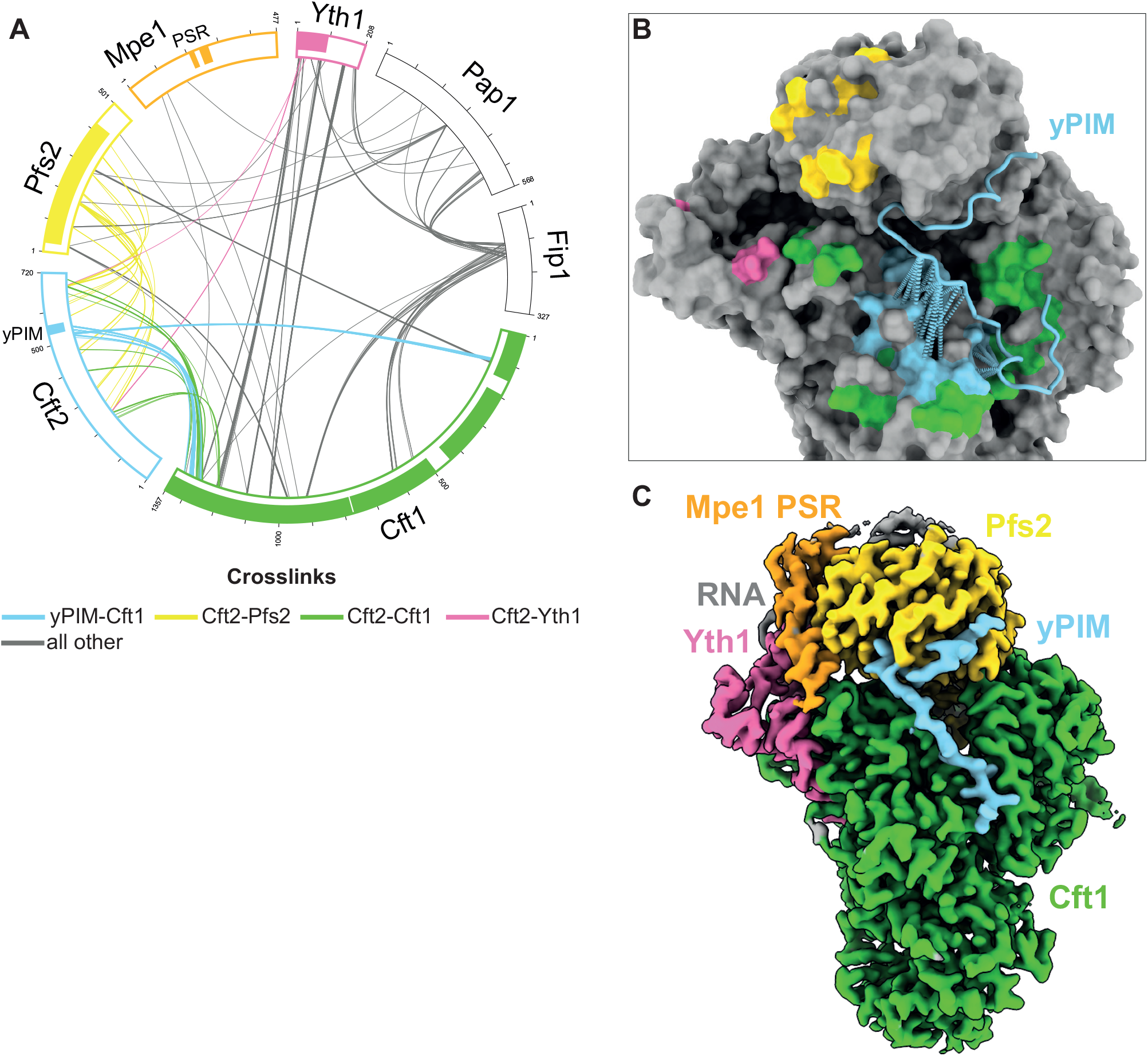
Cft2 antagonizes Mpe1 binding to polymerase module. (A) Circular view of crosslinking mass spectrometry analysis of a polymerase module- Cft2(S)-Mpe1-RNA complex. Each line represents a crosslinked peptide between the two proteins it connects. Colored lines correspond to Cft2(S) crosslinks with a subunit of polymerase module as indicated. White boxes represent the entire length of the corresponding protein, and boxes of solid color represent regions that are visible in the polymerase module-Mpe1-RNA and polymerase module-Mpe1-Cft2(S)-RNA structures. (B) Surface representation of the polymerase module-Mpe1-Cft2(S)-RNA structure (grey) highlighting regions where Cft2(S) crosslinks to Pfs2 (yellow), Yth1 (pink) and Cft1 (green). Crosslinks between the yPIM and Cft1 are shows as pseudobonds (light blue dotted lines) and light blue surfaces on Cft1. (C) CryoEM map of polymerase module-Mpe1-yPIM-RNA complex. The sample for this complex contains polymerase module, Mpe1, a yPIM peptide from Cft2, and the pre-cleaved *CYC1* RNA. See also Figure S4.

We next determined the impact of an isolated yPIM peptide on complex formation. Size exclusion chromatography showed that substoichiometric amounts of Mpe1 co-migrate with the polymerase module in complex with a synthetic yPIM peptide (Figure S4A, top gel). Including the pre-cleaved *CYC1* RNA stabilizes Mpe1 on the complex (Figure S4A, middle and bottom gels). In agreement with this, we were able to obtain a cryoEM map of this complex to a resolution of 2.6 Å that shows that both the Mpe1 PSR and yPIM can bind to the polymerase module simultaneously in the presence of RNA (Figures 4C and S4B-E).

These data suggest that an RNA-dependent structural rearrangement allows Mpe1 to stably bind to the polymerase module but the position of Cft2 on the polymerase module may sterically hinder Mpe1 binding in the absence of RNA. Alternate docking of Mpe1-RNA and Cft2 on the polymerase module may be indicative of different functional states of CPF.

### Mpe1 activates CPF cleavage and polyadenylation activity

To determine the functional role of Mpe1 in mRNA 3ʹ-end processing we purified a fully recombinant 14-subunit CPF with and without Mpe1 (CPF^ΔMpe1^) (Kumar et al., 2021) (Figure S5A-B). Both complexes were stable in size exclusion chromatography and showed correct stoichiometry of all subunits (Figure 5A).

**Figure 5.**
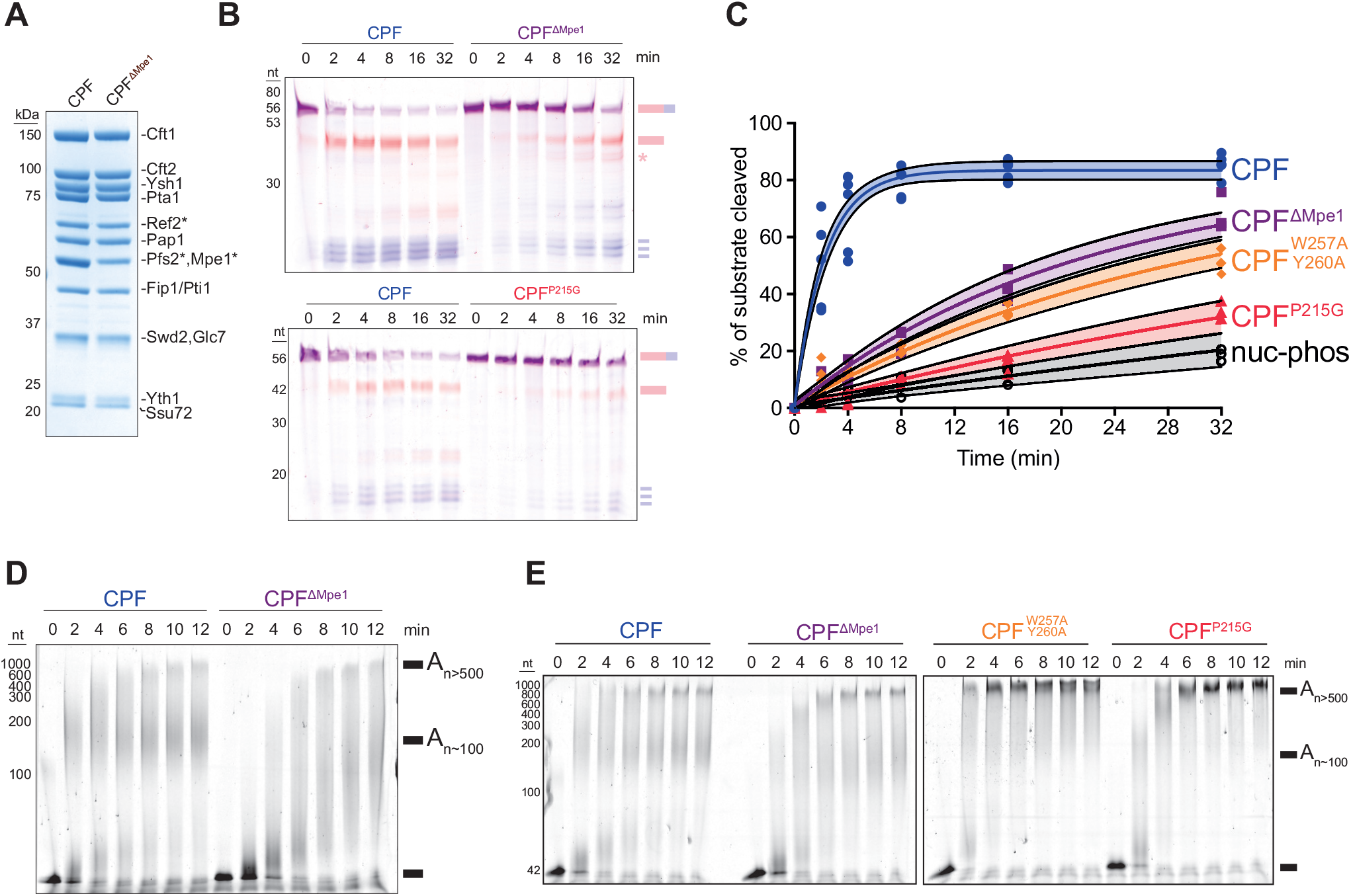
Mpe1 is a major regulator of CPF cleavage and polyadenylation activity. (A) SDS-PAGE of 10 pmol of CPF with (CPF) and without Mpe1 (CPF^ΔMpe1^). Subunits with an asterisk (*) denote they are SII-tagged. (B) Representative urea-PAGE of a dual-color *in vitro* cleavage assay using an uncleaved *CYC1* RNA substrate (5ʹ FAM and 3ʹ Alexa647 labels), and CPF, CPF^ΔMpe1^ or CPF^P215G^ as indicated. Cartoons of the substrate and expected RNA products are shown at the right. The asterisk indicates alternative 5ʹ cleavage products. (C) Quantitation of cleavage assays (as % of substrate cleaved) using CPF, CPF^ΔMpe1^ or CPF with mutant Mpe1, as indicated. Replicate data was fitted using global non-linear regression with an exponential plateau function and constraining the Ymax and Y0 values across complexes. For each complex the non-linear regression is shown as a solid line, and the shaded envelope represents the 95% confidence interval of the fit. Values for individual replicates are n = 5 for CPF, and n = 3 for all others. ‘nucl-phos’ is the CPF lacking the polymerase module. (D) Urea-PAGE of *in vitro* polyadenylation assay using a 5ʹ FAM-labeled pre-cleaved *CYC1* RNA substrate. Reactions were carried out with either CPF or CPF^ΔMpe1^ as indicated, using 100 nM CF IA and IB. (E) Similar to (D) except that CPF with mutant Mpe1 (CPF^W257A/Y260A^ and CPF^P215G^) were included as indicated, and reactions were carried out using 450 nM CF IA and IB. Note that in polyadenylation assays, the fluorescent signal is proportional to the amount of RNA regardless of its length. See also Figure S5.

We next performed *in vitro* cleavage assays with CPF, CF IA and CF IB using a dual-labelled fluorescent ‘uncleaved’ *CYC1* RNA substrate that includes the cleavage site (Figure 2A) (Hill et al., 2019). We monitored the cleavage reaction by resolving both the 5ʹ (FAM-labelled) and 3ʹ (Alexa647-labelled) cleavage products on a denaturing polyacrylamide gel. Cleavage assays showed that CPF^ΔMpe1^ is ∼10 times slower than full CPF (CPF t50 = 1.5 ± 0.27 min, CPF^ΔMpe1^ t50 = 14.9 ± 2.5 min; t50 is the time needed to cleave half of the maximum RNA cleaved by CPF; 83.4%) (Figure 5B-C). This suggests that Mpe1 is a major activator of CPF cleavage activity. Interestingly, there is increased cleavage at alternative sites for CPF^ΔMpe1^ (Figure 5B, asterisk), consistent with a role for Mpe1 in contacting the PAS and selecting the cleavage site. A CPF complex lacking the polymerase module did not show any cleavage activity (Figure S5C-D), confirming the essential role of the polymerase module in 3ʹ-end processing.

To test whether Mpe1 also stimulates polyadenylation activity in the context of full CPF, CF IA and CF IB, we performed *in vitro* polyadenylation assays using the 5ʹ FAM-labelled pre-cleaved *CYC1* substrate. Compared to the full complex, CPF lacking Mpe1 shows slower polyadenylation activity, and appears to be more distributive (Figure 5D). This suggests that Mpe1 plays an important role in promoting efficient polyadenylation. Thus, Mpe1 functions in both activating cleavage and controlling polyadenylation by CPF.

### Mpe1 interaction with Ysh1 is important for promoting cleavage and polyadenylation

We have shown that Mpe1 associates with CPF via two mechanisms: 1) through an interaction with Ysh1 (Hill et al., 2019), and 2) in the presence of RNA, Mpe1 rearranges to bind the polymerase module and the PAS. To test the importance of the interaction between the Mpe1 UBL domain and Ysh1, we purified a variant of Mpe1 containing four point mutations in conserved residues at the UBL-Ysh1 interface (F9A, D45K, R76E, P78G; Mpe1^FDRP^ henceforth). The Mpe1^FDRP^ variant did not assemble into CPF (Figure S5E). In contrast, Mpe1^FDRP^ co-eluted with the polymerase module in size exclusion chromatography (Figure S5F), comparable to its wild-type counterpart (Figure S1B). Thus, because Cft2 antagonizes the interaction between Mpe1 and the polymerase module, Mpe1 can no longer incorporate into CPF when its interaction with Ysh1 is disrupted. Mpe1^FDRP^ can still associate with the polymerase module in the absence of Cft2.

Next, we reasoned that although Mpe1^FDRP^ did not form a stable complex with CPF, its interaction with polymerase module might be sufficient to activate cleavage. To test this possibility, we used the purified CPF^ΔMpe1^ complex in the dual-color *in vitro* cleavage assay, and added 4x molar excess of Mpe1 in *trans*. Addition of WT Mpe1 activated CPF cleavage activity (Figure S5G). However, when we added Mpe1^FDRP^, we did not observe any stimulation of cleavage activity. This result suggests that the Mpe1- Ysh1 interaction is essential, not just for tethering Mpe1 to CPF, but also for activation of its endonuclease activity.

We also tested whether addition of WT or Mpe1^FDRP^ could stimulate the polyadenylation activity of CPF. We found that WT Mpe1 fully restored CPF polyadenylation activity, but Mpe1^FDRP^ could only partially rescue it (Figure S5H). Together, these data are consistent with a role for the interaction of the Mpe1 UBL with Ysh1 in stably tethering Mpe1 to CPF to promote both cleavage and regulated polyadenylation.

### Mpe1 PSR is required for cleavage and regulated polyadenylation

To test the functional relevance of the Mpe1 PSR, we generated mutants of Mpe1 that would disrupt its interaction with Pfs2 (W257A/Y260A) or its direct contact with A2 of the PAS (P215G). Both mutants could be incorporated into recombinant CPF (Figure S5I), in further agreement that Mpe1 is incorporated into CPF primarily through its interaction with Ysh1.

We next tested the cleavage and polyadenylation activities of CPF complexes carrying each of the Mpe1 PSR mutants. Both mutant complexes show dramatically reduced endonuclease activity compared to WT CPF or CPF^ΔMpe1^ (CPF ^W257A/Y260A^ t50 = 21.1 ± 3.9 min, CPF^P215G^ t50 = 46.3 ± 12.1 min) (Figure 5B-C). These data suggest that PSR interaction with polymerase module and RNA is crucial for activation of cleavage.

Surprisingly, in polyadenylation reactions, both PSR mutant complexes show aberrant hyperpolyadenylation compared to WT or CPF^ΔMpe1^ complexes (Figure 5E). Thus, when the PSR is disrupted, polyadenylation is deregulated.

Together, these data suggest that Mpe1 must interact with Ysh1, Pfs2 and RNA, for its ability to fully activate CPF cleavage, and regulate polyadenylation activity. Interestingly, Mpe1 PSR mutants cause more severe defects in both cleavage and polyadenylation than complete loss of Mpe1, suggesting that the presence of a defective Mpe1 might prevent other subunits of CPF, CF IA or CF IB from compensating for lack of Mpe1.

### Mpe1 is required for timely transcription termination

The latent cleavage activity of CPF^ΔMpe1^ raised the possibility that CPF may still cleave nascent RNA and commit RNAPII to termination in cells, even in the absence of Mpe1. To address this possibility, we investigated the consequence of acute Mpe1 depletion on the yeast transcriptome. We inserted a mini-auxin induced degron (mAID) (Tanaka et al., 2015) at the C-terminal end of the endogenous *MPE1* locus (Mpe1- mAID). Depletion of Mpe1 occurs within 30 min of adding 1 mM auxin to Mpe1-mAID cells and results in dose-dependent growth arrest (Figure S6A-B). To circumvent buffering mechanisms that could potentially mask the immediate impact of Mpe1 depletion on the transcriptome (Haimovich et al., 2013; Rodriguez-Molina et al., 2016; Sun et al., 2013), we isolated nascent transcripts by labelling newly-made RNAs with 4- thiouracil, followed by their biotinylation, purification and analysis (Sun et al., 2012) (Figure S6C). We also included a 4-thiouracil-labelled *S. pombe* spike-in as an independent internal control.

Mpe1 depletion followed by RT-qPCR of the nascent and total RNA revealed that Mpe1 depletion primarily impacted mRNA genes with very little impact on small nucleolar RNA genes (Figure S6D). This is consistent with a gene-class specific function of CPF in 3ʹ-end processing of mRNA genes (Lidschreiber et al., 2018).

Together these data show that we are able to rapidly and acutely deplete Mpe1 from yeast cells with an impact on mRNA transcription, consistent with Mpe1 function.

We next sequenced untreated or auxin-treated total and nascent RNA fractions from WT and Mpe1-mAID cells. Auxin treatment caused a significant change in the nascent RNA levels of 2,623 genes in Mpe1-mAID cells (1,459 increased, 1,164 decreased, log2-fold change >0, FDR-adjusted p-value <0.05), but had no significant impact on transcripts in WT cells (Figure S6E). Overall, depletion of Mpe1 showed a stronger impact on the nascent RNA fraction compared to total RNA (change in 1,661 genes, 866 increased, 795 decreased, Figure S6E). We therefore focused subsequent analyses on the nascent fraction which would more faithfully reflect the immediate impact of Mpe1 depletion.

Mpe1 depletion led to a widespread increase in the signal downstream of the polyadenylation site of mRNA genes (Figure 6A-B, red). Since the nascent RNA downstream of the cleavage site is normally rapidly degraded after CPF-mediated cleavage of the pre-mRNA, our observation is consistent with defects both in 3ʹ-end processing and transcription termination, and is indicative of RNAPII readthrough beyond the normal 3ʹ end of the transcript. The termination defect upon depletion of Mpe1 was similar to the strong defect observed upon nuclear depletion of Ysh1 (Baejen et al., 2017) (Figure 6B, dark grey). Thus, the latent endonuclease activity of CPF^ΔMpe1^ that we observe *in vitro* is largely insufficient to promote timely RNAPII termination *in vivo*.

**Figure 6.**
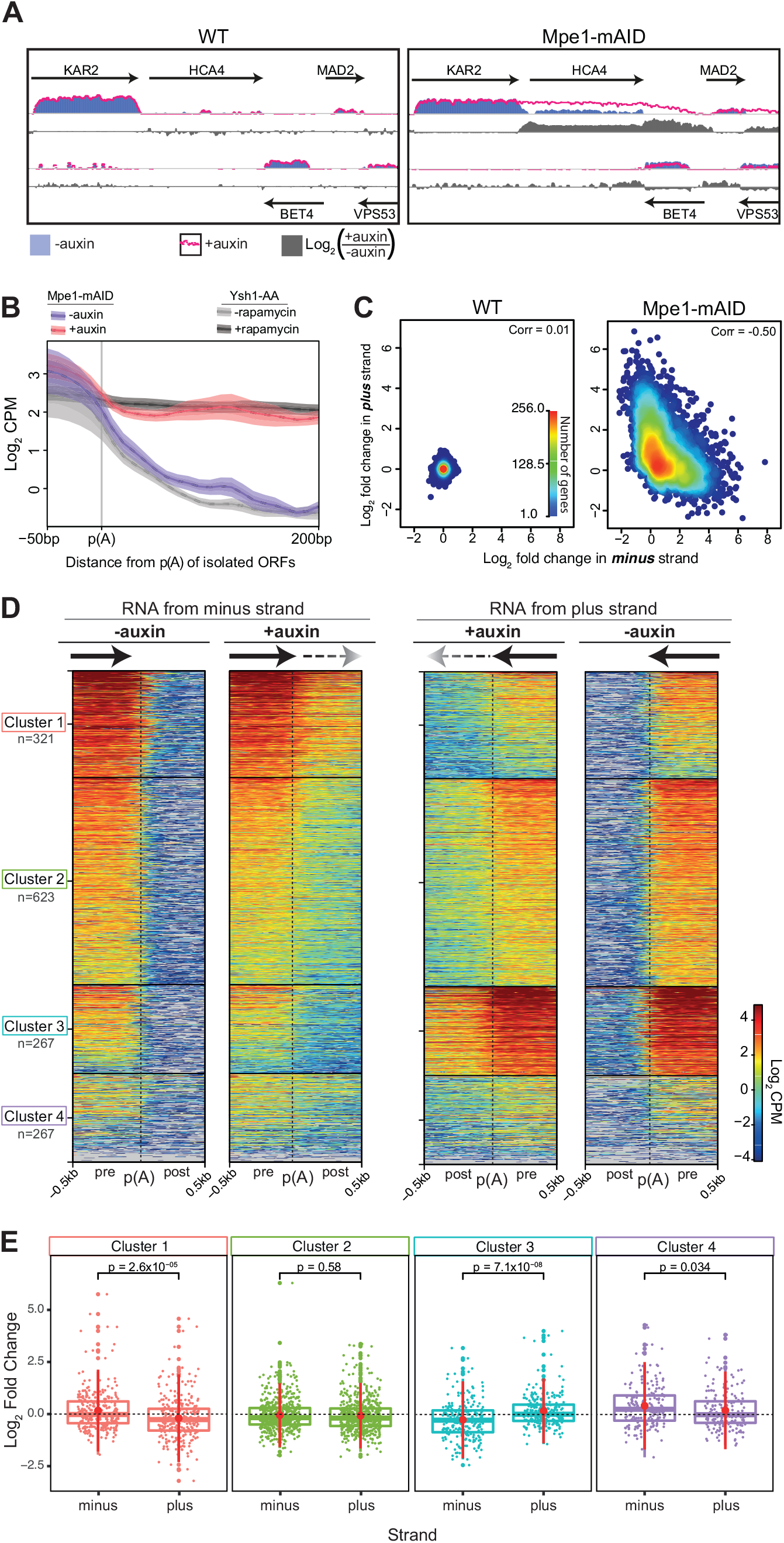
Mpe1 is globally required for timely transcription termination. (A) Representative genomic snapshot of overlaid profiles of strand-specific nascent RNAseq from WT or Mpe1-mAID yeast cells either untreated (-auxin, blue bars) or treated with 1 mM auxin for 30 min (+auxin, magenta trace). The log2 fold change in nascent RNA upon addition of auxin is displayed as grey bars. Data are shown for each strand as indicated by the directionality of arrows, which represent protein-coding genes. (B) Metagene plots of nascent RNA at the polyadenylation site (p(A)) from Mpe1-mAID cells treated with auxin (+auxin, magenta trace) or untreated (-auxin, blue trace). Nascent RNA from Ysh1 anchor away cells (Ysh1-AA), where Ysh1 was depleted (+rapamycin, dark gray) or not depleted from the nucleus (-rapamycin, light gray) is also shown (Baejen et al., 2017). Selected genes are at least 200 bp from neighboring ORFs (n = 931). (C) Density scatter plot of changes in nascent RNA synthesis in WT or Mpe1-mAID cells upon addition of auxin. Values correspond to strand-specific log2 fold change per gene vs. the corresponding calculation on the opposite strand. (D) k-means-clustering of strand-specific nascent RNA before (-auxin) and after auxin treatment (+auxin) in Mpe1-mAID cells. Data are shown for a 1 kb window centered around the poly(A) site (p(A), vertical dotted line) of 1,478 convergent gene pairs. For clustering, *k* = 4 was determined empirically to give clusters with distinct patterns. Transcription directionality is indicated with arrows. The number of genes in each cluster (n) is indicated. Log2 CPM: log2 of the counts per million. (E) Log2 fold change in nascent RNA upon Mpe1 depletion for the genes in each of the clusters in (D). Each boxplot corresponds to the group of genes in the indicated orientation (minus or plus strand). Dots represent the log2 fold change for each gene (calculated with DESeq2), large dots represent outliers within each distribution. P-values are from pairwise Student’s t-test. Middle horizontal line in each boxplot represents the median, top and bottom lines of each box show the interquartile range. Dashed line indicates no change. See also Figure S6.

Closer inspection of the data revealed that the magnitude and apparent length of readthrough transcription varied depending on the local orientation of genes. A termination defect in *KAR2*, for example, produced readthrough transcripts that appeared to invade the downstream co-directional *HCA4* gene and terminated within the next gene, *BET4*, located on the opposite strand (Figure 6A). To investigate whether there was a relationship between changes in nascent RNA signal and transcription orientation, we calculated the strand-specific log2-fold change in the nascent RNA signal for every gene across the genome, and compared it to the change in RNA signal on the corresponding position in the opposite strand. We also performed a similar analysis using a sliding 1 kb window across the genome. In both cases, the changes in RNA synthesis upon Mpe1 depletion are anti-correlated between strands (Figures 6C S6F).

These analyses suggest that readthrough transcription negatively impacts ongoing transcription on the opposite strand, and is consistent with transcription interference (Shearwin et al., 2005).

To specifically analyze transcription interference, we focused our subsequent analyses on the impact that readthrough transcription has on pairs of genes. We carried out a *χ*^2^ test of independence between the impact of Mpe1 depletion on nascent RNA and gene organization (Figure S6G). This analysis revealed a positive association between increased nascent RNA levels and co-directionally oriented pairs of genes.

This association is likely due to readthrough defects increasing the nascent RNA signal of the downstream gene in a co-directional pair (*i.e. HCA4*; Figure 6A). Decrease in nascent RNA on the other hand was associated with genes that share both a convergent and divergent partner (*i.e.* a gene with both a divergent upstream and a convergent downstream partner gene) (Figure S6G). This result is consistent with convergently-oriented pairs of genes having an antagonistic impact on each other upon Mpe1 depletion.

To test if readthrough transcription was uniform across the genome, we performed k-means clustering of the strand-specific nascent RNA signal at the 3ʹ-end of convergent gene pairs (Achar et al., 2020). (Figure 6D). Our analyses reveal transcription readthrough beyond the poly(A) site in all clusters on both strands (minus and plus) in Mpe1-depleted cells (+auxin). The level of readthrough is proportional to the signal preceding the poly(A) site, indicating that the role of Mpe1 is independent of the baseline level of nascent RNA synthesis, and is thus globally required for transcription termination.

Finally, we explored whether transcription of either gene within a convergent pair is differentially impacted upon depletion of Mpe1. To that end we compared the log2-fold change between convergent gene pairs within each cluster (Figure 6E). This shows that genes with lower baseline transcription levels were more likely to show a decrease in nascent RNA than their counterpart (Figure 6E; plus strand in cluster 1, minus strand in cluster 3). In contrast, there is an equal impact when baseline RNA synthesis is comparable between both genes in a convergent pair (cluster 2). Given that the level of readthrough is correlated to the baseline level of RNA synthesis, the differential impact likely reflects transcription interference and RNAPII collision events that are skewed against genes with lower baseline transcription levels.

Our genomic analyses reveal that Mpe1 plays a global and essential role in the timely activation of CPF cleavage activity. Thus, CPF is required for transcription termination by RNAPII and is crucial for preventing transcription interference of neighboring genes.

## Discussion

The 3ʹ-end processing machinery couples recognition of conserved sequence elements in the nascent pre-mRNA with cleavage and polyadenylation. Here, we reveal that binding of the polyadenylation signal by the polymerase module is sensed by Mpe1 to promote efficient cleavage and polyadenylation. Specifically, we show: (1) Mpe1 is remodeled and stably binds to Pfs2 in the presence of RNA. This is required for endonuclease activation and regulated polyadenylation. (2) Cft2 antagonizes docking of Mpe1 onto polymerase module-RNA. Mpe1 and Cft2 both provide a direct link between the nuclease and polymerase modules. (3) An Mpe1-Ysh1 interaction stably tethers Mpe1 on CPF in the absence of RNA, and is essential for Ysh1 activation. (4) Mpe1 is essential for timely transcription termination across the genome.

### PAS recognition by Yth1 and Mpe1

RNA elements required for cleavage and polyadenylation are more degenerate in yeast than in humans (Guo and Sherman, 1996; Russo et al., 1993). Computational analyses show that the PAS is conserved as an A-rich sequence in yeast, frequently AAUAAA (Graber et al., 1999; Guo and Sherman, 1996; Russo et al., 1993; Tian and Graber, 2012). Alongside previous work (Clerici et al., 2018; Sun et al., 2018), our structure reveals that binding of Yth1 zinc finger 2 to the A1A2 dinucleotide is highly conserved across eukaryotes. Zinc finger 3 is expected to recognize the A3A4 dinucleotide but has not yet been observed in any cryoEM maps of the yeast complex, possibly because it is flexible. We show that RNA binding may additionally involve one nucleotide upstream of the PAS since a U is bound in a pocket on the surface of Yth1, suggesting that the sequence context of the PAS is important.

In the Mpe1-bound structure, the A2 base is additionally contacted directly by the conserved P215 in Mpe1 through a CH-π interaction. CH-π interactions are a special type of relatively weak hydrogen bond between a partially charged proton and the delocalized electron π system of an aromatic group (Chakrabarti and Samanta, 1995; Nishio et al., 2014). In principle, CH-π interactions can involve protons from almost any amino acid. Thus, the conservation of a proline at this position may indicate that it is playing a dual role; (i) as a general sensor of RNA binding by Yth1 through a CH-π interaction, and (ii) to enforce the correct fold of the Mpe1 PSR due to its sterically restricted side chain. The P215-RNA contact appears to stabilize the PSR of Mpe1 on polymerase module and is likely to be essential in sensing the RNA bound state of CPF.

### An interplay between Mpe1 and Cft2 regulates CPF activation

Cft2, in addition to its function in tethering the nuclease module to the polymerase module via the yPIM, appears to play a role in regulating Mpe1 docking onto the polymerase module: when Cft2(S) was included in our cryoEM sample, Mpe1 and RNA were no longer visible. This suggests that Cft2 antagonizes Mpe1 PSR binding to the polymerase module. In agreement with this, multiple regions of Cft2 are located in close proximity to the Mpe1 PSR binding site (Figure 4B) and may block its binding.

Interestingly, even though they are not visible, RNA and Mpe1 remain bound to the polymerase module-Cft2 complex. Thus, it is likely that there are additional interactions between the polymerase module and Mpe1 and/or RNA that are not visible in the cryoEM maps, possibly because they are dynamic. Moreover, it remains unclear whether other CPF subunits stabilize simultaneous direct Cft2 yPIM and Mpe1 PSR binding to the polymerase module in the context of CPF. Together, the dynamics of Cft2 and Mpe1 may help define distinct functional states of CPF that both safeguard and promote activation of Ysh1.

### Efficient CPF activation is essential for safeguarding the transcriptome

Our transcriptomic studies suggest that Mpe1 is globally required for efficient activation of cleavage activity and, consequently, timely transcription termination. In the absence of Mpe1, there is extensive readthrough and transcription interference on downstream genes. CPF activity may be particularly important in yeast for preventing transcription interference: head-to-head RNAPII collisions occur at a basal level between convergent genes under normal circumstances (Hobson et al., 2012), suggesting that any delay in termination is deleterious. Moreover, because intergenic regions in the yeast genome are relatively short compared to mammalian genomes (Chen et al., 2012; Xu et al., 2009), CPF must be activated as soon as the PAS is recognized. In human cells, closely-arranged convergent genes show transcription interference upon depletion of the CPSF endonuclease CPSF73 (Eaton et al., 2020), and the mutations in the β-globin 3ʹ-UTR that cause β-thalassemia lead to termination defects and transcription interference on the downstream gene (Proudfoot, 1986; Whitelaw and Proudfoot, 1986). Thus, in addition to 3ʹ-end processing, CPF/CPSF universally safeguards the transcriptome from unwanted transcription interference.

### RBBP6 may also activate cleavage in human CPSF

The human homolog of Mpe1, RBBP6, is known to play a role in 3ʹ-end processing and has a similar domain organization to Mpe1 (UBL, zinc knuckle and RING finger) with a metazoan-specific C-terminal extension containing p53 and retinoblastoma binding sites (Di Giammartino et al., 2014; Pugh et al., 2006). Given the high degree of conservation between yeast Mpe1 and human RBBP6 in the PSR loop that contacts RNA, it is likely that a similar RNA-sensing mechanism operates across eukaryotes. In the human mPSF, however, CPSF30 occupies the hydrophobic pocket on WDR33 where the RBBP6 helix would bind (Zhang et al., 2020). It therefore seems likely that RBBP6 senses RNA binding in the same way as Mpe1, but RBBP6 interaction with WDR33 may differ.

### Model for CPF activation cycle

Based on these studies, we propose a new model for CPF activation (Figure 7): Prior to RNA binding, Mpe1 is flexibly tethered to CPF, primarily through its interaction with Ysh1. Cft2 impairs stable docking of Mpe1 onto the polymerase module, which keeps Ysh1 in an inactive state, thereby avoiding precocious activation of cleavage. Studies of yeast CPF and human CPSF suggest that in this configuration, the endonuclease is too far from the cleavage site and requires a spatial rearrangement to reach its substrate (Hill et al., 2019; Zhang et al., 2020).

**Figure 7.**
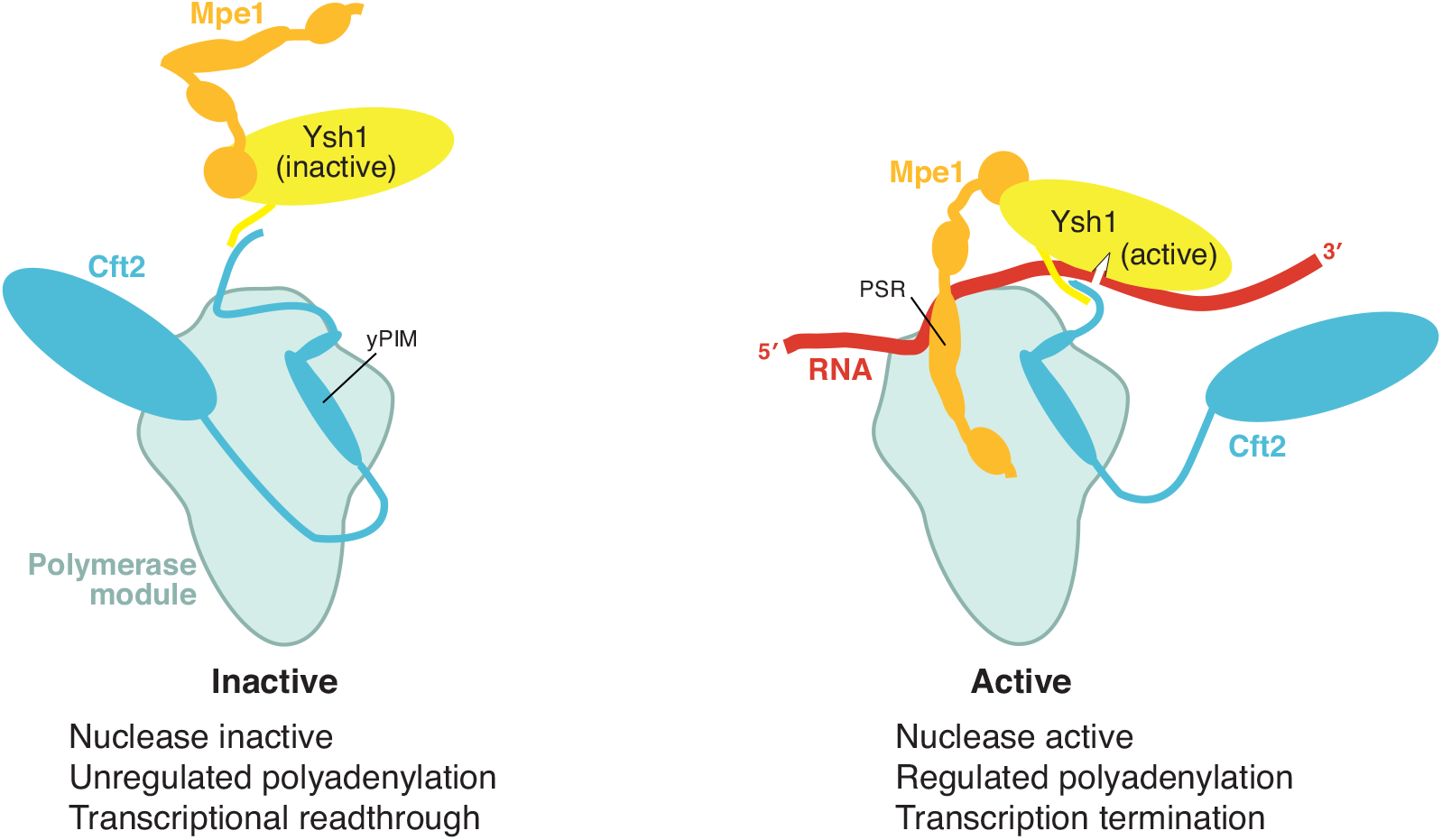
Mpe1 activates CPF cleavage and regulates polyadenylation. In the inactive state (left), Mpe1 is flexibly tethered to the complex via Ysh1. Cft2 tethers the nuclease module to the polymerase module through its yPIM, and restricts Mpe1 association with the polymerase module. RNA binding, CF IA and CF IB are required for activation of CPF (right). Recognition of the PAS by Yth1 in the polymerase module triggers an Mpe1-driven rearrangement of the nuclease module that may bring Ysh1 in close proximity to the cleavage site.

When Yth1 recognizes a newly-transcribed polyadenylation signal, Mpe1 senses the bound RNA through its PSR and docks onto the polymerase module. This RNA-induced remodeling of Mpe1, coupled with the direct interaction between Mpe1 and Ysh1, repositions Ysh1 in close proximity to the cleavage site on RNA. In this RNA-bound state, Mpe1 may license the assembly of CPF, CF IA, CFIB and RNA into a configuration that is activated for cleavage. Previous studies suggested that Mpe1 and the polymerase module directly interact with CF IA (Casanal et al., 2017; Lee and Moore, 2014; Vo et al., 2001), which is required for activation of cleavage (Hill et al., 2019). Therefore, one possibility is that docking of Mpe1 onto the polymerase module forms a new interaction surface for CF IA to activate cleavage. Interestingly, RBBP6 also interacts with CstF64, the human homolog of the CF IA subunit, Rna15 (Di Giammartino et al., 2014). Taken together, Mpe1 and RBBP6 may play the same role to link CF IA/CstF, CPF/CPSF, and RNA binding to activation of the endonuclease. We also speculate that this remodeling could activate Glc7 in the phosphatase module to promote transcription termination (Schreieck et al., 2014).

Several factors ensure the fidelity of 3ʹ-cleavage and polyadenylation. Multiple mechanisms regulate transitions between active and inactive states to control mRNA 3ʹ- end processing, to prevent inappropriate activation of cleavage and to control the length of the poly(A) tail. We showed here that Mpe1 senses when the PAS sequence is bound to CPF, thereby enforcing the correct choice of polyadenylation site. Together, these data suggest that, similar to splicing, recognition and enzymatic processing of an RNA substrate governs CPF remodeling from an inactive to an active state to control mRNA processing.

## Acknowledgements

We thank Grigory Sharov, Ananthanarayanan Kumar, Shabih Shakeel, Manuel Carminati, Viswanathan Chandrasekaran, Eva Absmeier and Katerina Naydenova for assistance with cryoEM and image processing; Chris H. Hill for truncated Cft2; M. Cemre Manav for assistance with protein purification; Juan Mata for *S. pombe* strains; Jake Grimmett and Toby Darling (LMB scientific computation) for support; Jianguo Shi for assistance with baculovirus; and Ana Casañal, Alexey Murzin, and members of the Passmore lab for helpful discussions and advice. This work was supported by the Medical Research Council, as part of United Kingdom Research and Innovation, MRC file reference number MC_U105192715 (L.A.P.); the European Union’s Horizon 2020 research and innovation programme (ERC Consolidator grant agreement 725685) (to L.A.P). The Wellcome Centre for Cell Biology is supported by core funding from the Wellcome Trust [203149]. We acknowledge the MRC Laboratory of Molecular Biology Electron Microscopy Facility for access and support of electron microscopy sample preparation and data collection, and Andre Howe and Diamond Light Source for access to eBIC (proposal BI23268) funded by the Wellcome Trust, MRC and Biotechnology and Biological Sciences Research Council.

## Author Contributions

J.B.R.-M. performed experiments, analyzed data and wrote the manuscript. E.S. performed Mpe1 depletion studies and prepared RNA-seq libraries. S.M. and J.S.M. performed and analyzed the HDX-MS experiments. F.J.O. and J.R. performed and analyzed the crosslinking MS experiments. L.A.P. supervised the research, analyzed the data and wrote the manuscript. All authors contributed to the writing of the manuscript.

## Materials and Methods

### Cloning

Oligonucleotides and plasmids used in this study are listed in Tables S1 and S2, respectively.

#### Pairwise combinations of Mpe1 and polymerase module subunits

Twin strep (SII)-tagged Mpe1 was amplified from pIDS-Mpe1 (Hill et al., 2019) using primers pIDS_CasII_F and pIDS_Casω-R. *E. coli* codon-optimized genes encoding Cft1, Pap1, Pfs2, Fip1 and Yth1 (GeneArt) were amplified using primers pB/pIDC_CasI_F and pB/pIDC_CasI_R. Mpe1 and one of the polymerase module genes were cloned into a SwaI (NEB) digested pBig1a vector of a modified biGBac system (Hill et al., 2019; Weissmann et al., 2016) via Gibson assembly. Colonies were screened for correct constructs by restriction digest using BamHI and XbaI (NEB).

#### Polymerase module-Mpe1

pBig1a carrying genes encoding subunits of the polymerase module (Cft1, Pap1, Pfs2, Fip1 and Yth1) (Kumar et al., 2021) and pBig1b carrying Mpe1-3C-SII were digested with PmeI (NEB) to release the gene cassettes. Cassettes were cloned into PmeI digested pBig2ab via Gibson assembly and transformed into chemically competent Top10 *E. coli*. Constructs were screened via SwaI restriction digestion.

#### Mpe1 variants

The Mpe1^FDRP^ variant (F9A, D45K, R76E, P78G) was generated by Gibson assembly of PCR products using the corresponding primers listed in Table S1. PCR products have overlapping sequences to allow for Gibson assembly into the pMA vector. Mpe1^FDRP^ was subcloned via PCR and Gibson assembly into a modified pACEBAC1 vector that introduces an in-frame TEV cleavage site followed by an SII tag at the 3ʹ-end of the insert (pACEBAC_TEV_SII). Mpe1^P215G^ and Mpe1^W257A/Y260A^ variants were similarly generated with PCR products carrying the respective mutations and Gibson assembly into a BamHI (NEB) digested pACEBAC_TEV_SII vector. Constructs were screened and mutations confirmed by Sanger sequencing (Source Biosciences) using the pACE_Mpe1_F primer.

### Baculovirus-mediated insect cell protein overexpression

Preparation of bacmids in EMBacY cells, transfection into *Sf9* cells, initial and secondary baculovirus generation, and protein overexpression in *Sf9* cells were carried out as previously described (Hill et al., 2019; Kumar et al., 2021).

### Protein purification

Protein purification was carried out essentially as previously described (Casanal et al., 2017; Hill et al., 2019; Kumar et al., 2021) with several modifications as described below.

Polymerase module carrying an SII-tag on Pfs2 was purified as previously described (Casanal et al., 2017), except that RNase and DNase were omitted from the lysis buffer and 1 ml of BioLock (IBA, cat. No. 2-0205-050) was added.

For polymerase module-Mpe1, a frozen pellet of insect cells (3-6 L at 2x10^6^ cells/ml) expressing polymerase module and SII-tagged Mpe1 were lysed in Buffer A (50 mM HEPES pH 8, 75 mM NaCl, 1 mM TCEP) supplemented with 3x protease inhibitor tablets (Roche, cat. No. 11836153001) and 1 ml BioLock. Cleared lysate was applied to Strep-Tactin resin (IBA, cat. No. 2-1201-025) and washed with Buffer A. Protein was eluted with 20 ml of Buffer A supplemented with 1.2 mg/ml desthiobiotin (IBA, cat. No. 2-1000-005). Eluate was filtered through a 0.45 µm filter and applied to a 1 ml MonoQ 5/50 GL column (Cytiva, cat. No. 17516601) equilibrated in Buffer A. The complex was eluted off the column using a linear gradient up to 50% Buffer B (50 mM HEPES pH 8, 1 M NaCl, 1 mM TCEP) over 40 column volumes. During elution, Mpe1 and polymerase module dissociate, elute as distinct peaks, and were kept separate following purification. Individual fractions containing Mpe1 or polymerase module were pooled and concentrated using a 30 kDa or 100 kDa cut-off concentrator, respectively. Concentration was determined using a nanodrop (ThermoFisher) and the theoretical extinction coefficient for Mpe1 (32,430 M^-1^ cm^-1^) or polymerase module (298,530 M^-1^ cm^-1^) calculated in ProtParam (Gasteiger, 2005). The polymerase module from this purification is untagged. Purified protein was flash frozen in liquid N2 and kept at -80°C.

The polymerase module-Cft2 complex was purified from insect cells as described for the polymerase module-Mpe1 complex except that it was from cells co-infected with two viruses (in a 1:1 ratio); one carrying polymerase module (with an SII tag on Pfs2) and another carrying untagged Cft2 and Ysh1. Ysh1 did not stably co-purify with the rest of the complex. Concentration was determined using the theoretical extinction coefficient for the polymerase module-Cft2 complex (383,790 M^-1^ cm^-1^) calculated in ProtParam.

SII-tagged Mpe1 variants were purified following the same procedure described for polymerase module-Mpe1 complex with some modifications. Mpe1 was purified from 1-3 L at 2x10^6^ cells/ml of pelleted and frozen insect cells. The filtered eluate from the Strep-Tactin affinity purification was applied to a HiTrap Heparin 1ml column (Cytiva, cat. No. 17040601).

#### Cft2(S)

9L of BL21* cells carrying the pET28a +(modified) 6H-3C-Cft2(S) vector were induced with 0.5 mM IPTG overnight at 18°C. Cells were lysed in Buffer C (50 mM HEPES pH 7.9, 30mM imidazole, 250 mM NaCl, 1 mM TCEP) supplemented with protease inhibitor tablets, DNaseI and RNaseA. Cleared lysate was incubated with Ni-NTA (Qiagen) resin, washed with Buffer C and eluted with Buffer D (50 mM pH 7.9, 500 mM imidazole, 250 mM NaCl). Pooled fractions were cleaved with 3C protease (140 μg/ml) at 4°C. Cleaved Cft2(S) was further purified by anion exchange chromatography using a 1 ml MonoQ 5/50 GL column equilibrated in Buffer E (50 mM HEPES pH 7.9, 150 mM NaCl, 1 mM TCEP) over a gradient up to 100% over 100 column volumes using Buffer F (50 mM HEPES pH 7.9, 1 M NaCl, 1 mM TCEP). Pooled fractions containing Cft2(S) were concentrated using a VivaSpin concentrator (30 kDa cutoff) and further purified by size exclusion chromatography using a HiLoad 16/600 Superdex 200 pg column (Cytiva, 28989335) equilibrated in Buffer E. Fractions containing Cft2(S) were pooled and concentrated as before and flash frozen in liquid N2 and stored at - 80°C.

For the complex of Ysh1-Cft2-phosphatase module, insect cells were co-infected (1:1 volume ratio) with two viruses; one carrying genes encoding Ysh1 and Cft2 and another carrying genes encoding subunits of the phosphatase module with an SII tag on the Ref2 subunit. Cells were lysed in Buffer G (50 mM HEPES pH 8, 150 mM KCl, 0.5 mM Mg(OAc)2, 0.5 mM TCEP) supplemented with 3x protease inhibitor tablets and 1 ml BioLock. Resin was washed with Buffer C and eluted from the Strep-Tactin resin with 1.2 mg/ml desthiobiotin. Eluate was filtered through a 0.45 µm filter and applied to a 1 ml MonoQ 5/50 GL column equilibrated in Buffer C. The complex was eluted off the column using a linear gradient up to 50% Buffer H (50 mM HEPES pH 8, 1 M KCl, 0.5 mM Mg(OAc)_2_, 0.5 mM TCEP) over 40 column volumes. Fractions with eluted protein showing correct stoichiometry were pooled, concentrated in a VivaSpin concentrator (100 kDa cutoff) and flash frozen. Concentration was determined using the theoretical extinction coefficient for Ysh1-Cft2-phosphatase module (334,400 M^-1^ cm^-1^) calculated in ProtParam.

### *In vitro* reconstitution of CPF^ΔMpe1^

Purified protein complexes (5 µM polymerase module, 5 µM Ysh1-Cft2- phosphatase module) and 15 µM Mpe1 or any variants thereof were mixed and the final volume was brought to 50 µl with SEC 150 KCl buffer (20 mM HEPES pH 8, 150 mM KCl, 0.5 mM Mg(OAc)2, 0.5 mM TCEP). Sample was kept on ice before being loaded on a Superose 6 Increase 3.2/300 (Cytiva, cat. No. 29091598) equilibrated in SEC 150 KCl buffer. Fractions showing all components of the assembled complex with correct stoichiometry were pooled, concentrated in a VivaSpin concentrator (100 kDa cutoff) and flash frozen. Concentration was determined using the theoretical extinction coefficient for CPF^ΔMpe1^ (632,930 M^-1^ cm^-1^) or CPF (665,360 M^-1^ cm^-1^) calculated in ProtParam.

### In vitro reconstitution of polymerase module with Mpe1 and Cft2 (or variants)

Complexes were assembled and purified as described for CPF with some modifications. 5 µM of polymerase module was mixed with 15 µM Mpe1 and either 10 µM Cft2(S) or 720 µM of a synthetic yPIM peptide (GenScript). For full length Cft2, 5 µM of the polymerase module-Cft2 complex was mixed with 15 µM Mpe1. In all cases where RNA was included, 10 µM of 5ʹ-FAM labelled pre-cleaved *CYC1* substrate (IDT; Table S3) was added to the complex. The final volume of the assembly was brought to 50 µl with SEC 50 buffer (20 mM HEPES pH 8, 50 mM NaCl, 0.5 mM TCEP). Fractions showing all components of the assembled complex with correct stoichiometry were pooled, concentrated in a VivaSpin concentrator (100 kDa cutoff) and flash frozen.

Concentration was determined using the theoretical extinction coefficient for polymerase module-Mpe1 (330,960 M^-1^ cm^-1^) calculated in ProtParam. For cryoEM sample preparation, 3 µl from the peak fraction were used per grid.

### CryoEM

#### Sample preparation and data collection

1.2/1.3 UltrAuFoil (Quantifoil) grids (Russo and Passmore, 2014) were glow discharged using an Edwards Sputter Coater S150B at setting 8 for 90 sec. Complexes were freshly assembled as described above, and 3 µl from the peak fraction was used per grid. Grids were frozen in liquid ethane using Vitrobot Mark IV (ThermoFisher) with 5 sec blotting, blotting force -10 at 4°C in 100% humidity.

For the polymerase module-Mpe1-RNA complex 12,060 movies were collected on Krios II at eBIC with a K3 detector in counting mode (bin 1), pixel size of 0.83 Å/pixel, total dose of 40 e^-^/Å^2^ with a defocus range of -0.5 µm to -3.1 µm in 0.2 µm steps.

For the polymerase module-Mpe1-Cft2(S)-RNA complex 5,118 movies were collected on Krios III at MRC-LMB with a K3 detector in counting mode (bin 1), pixel size of 0.86 Å/pixel, total dose of 36.9 e^-^/Å^2^ with a defocus range of -0.5 µm to -3.1 µm in 0.2 µm steps.

For the polymerase module-Mpe1-yPIM-RNA complex 19,524 movies were collected on Krios III at MRC-LMB with a K3 detector in super-resolution counting mode (bin 2), pixel size of 0.86 Å/pixel, total dose of 40 e^-^/Å^2^ with a defocus range of -0.5 µm to -3.1 µm in 0.2 µm steps.

#### CryoEM data processing

A general description of cryoEM data processing is provided below. For complex-specific details regarding the polymerase module-Mpe1-RNA complex, the polymerase module-Mpe1-Cft2(S)-RNA complex or the polymerase module-Mpe1-yPIM-RNA complex please refer to Figure S1F, Figure S3C or Figure S4B, respectively.

Multi-frame movies from each data collection were processed using Relion 3.1 (Zivanov et al., 2018). Per-micrograph beam-induced motion was estimated and corrected using MotionCor2 using a 5x5 grid (Zheng et al., 2017), and the CTF was estimated using Gctf (Zhang, 2016). Before further processing, the best micrographs were selected, first based on their estimated resolution (at least 5 Å), and then according to their figure of merit (at least 0.05). We next used the previously-reported polymerase module map (Casanal et al., 2017), low-pass filtered to 35 Å, for template-based particle picking. Particles were first extracted binned ∼5x to the pixel size indicated in the supplementary figure schematics. Depending on the number of particles, they were randomly split into four equally-sized groups and eventually re- grouped. 2D Classification was carried out for each group, and class averages without clear presence of particles were discarded. 2D classification was repeated as indicated. Next we used the aforementioned polymerase module map to carry out 3D classification with or without a mask. Classes with isotropic maps and distinct internal features were selected and 3D refined. Refined maps were 3D classified without image alignment and classes selected again based on map isotropy and clear internal features. Particles were then re-extracted to their original pixel size as indicated and 3D refined.

#### Model construction, refinement and analysis

Mpe1 and RNA were manually modelled into their respective densities using Coot [v. 0.9.5.1-pre] (Emsley and Cowtan, 2004; Emsley et al., 2010). The yPIM was modelled using the PIM of CPSF100 (Zhang et al., 2020) as an initial model and modified in Coot. Models were refined in Phenix Real Space Refine [v. 1.19.2-4158] (Adams et al., 2010) and Coot. Models and maps were further visualized and analyzed in ChimeraX [v. 1.2] (Goddard et al., 2018). Buried surface area was calculated using PDBePISA (Krissinel and Henrick, 2007).

Protein sequences for Cft1, Pfs2, Mpe1 and Yth1 from *Saccharomyces cerevisiae*, *Schizosaccharomyces pombe*, *Danio rerio*, *Homo sapiens*, *Mus musculus*, *Caenorhabditis elegans* and *Drosophila melanogaster* were aligned using ClustalW (Goujon et al., 2010; Sievers et al., 2011). The alignment was used to color the surface of the polymerase module-Mpe1-RNA structure using the entropy-based conservation index of AL2CO (Pei and Grishin, 2001) implemented in ChimeraX. Orientation distribution plots and Efficiency of orientation distribution (EOD) were calculated using cryoEF (Naydenova and Russo, 2017).

### Crosslinking mass spectrometry

The crosslinker sulfo-SDA (sulfosuccinimidyl 4,4′-azipentanoate) (Thermo Scientific Pierce) was dissolved in crosslinking buffer (20 mM HEPES, 50 mM NaCl, 0.5 mM TCEP, pH 8.0) to 4 mM before use. For photo-activation, the polymerase module-Cft2(S)-Mpe1-RNA complex (at 0.5 mg/ml in 250 µl) was incubated with sulfo-SDA at a final concentration of 2 mM for 2 h on ice. The sample was irradiated with UV light at 365 nm for 15 min. Crosslinked complex was visualized in Tris-Acetate gels (Thermo, cat. No. EA0375BOX). Gel slices containing the crosslinked complex were cut out with a clean razor blade and dehydrated with LCMS-grade acetonitrile. Samples were subsequently denatured in 8 M urea, 100 mM NH4HCO3. The proteins were derivatized with iodoacetamide and digested with LysC endoproteinase (Wako) for 4 h at 25 °C. After dilution of the sample to a urea concentration of 1.5 M with 100 mM NH4HCO3, trypsin (Thermo Scientific Pierce) was added and the samples were digested for 16 h at 25 °C. The resulting tryptic peptides were extracted and desalted using C18 StageTips.

Eluted peptides were fractionated using a Superdex 30 Increase 3.2/300 column (GE Healthcare) at a flow rate of 10 µl/min using 30 % (v/v) acetonitrile and 0.1 % (v/v) trifluoroacetic acid as mobile phase. 50 µl fractions were collected and dried. Five highest molecular weight samples for analysis were resuspended in 0.1 % (v/v) formic acid, 1.6 % (v/v) acetonitrile for LC-MS/MS. LC-MS/MS analysis was performed on an Orbitrap Fusion Lumos Tribrid mass spectrometer (Thermo Fisher) coupled on-line with an Ultimate 3000 RSLCnano system (Dionex, Thermo Fisher). Samples were separated on a 50-cm EASY-Spray column (Thermo Fisher). Mobile phase A consisted of 0.1 % (v/v) formic acid and mobile phase B of 80 % (v/v) acetonitrile with 0.1 % (v/v) formic acid. Flow rates were 0.3 μl/min using gradients optimized for each chromatographic fraction from offline fractionation, ranging from 2 % to 45 % mobile phase B over 90 min. Mass spec data were acquired in data-dependent mode using the top-speed setting with a three second cycle time. For every cycle, the full scan mass spectrum was recorded using the Orbitrap at a resolution of 120,000 in the range of 400 to 1,600 m/z. Ions with a precursor charge state between 3+ and 7+ were isolated and fragmented.

Each precursor was fragmentated by higher-energy Collisional Dissociation (HCD) at 26 %, 28 % and 30 %. The fragmentation spectra were then recorded in the Orbitrap with a resolution of 60,000. Dynamic exclusion was enabled with single repeat count and 60- seconds exclusion duration. A recalibration of the precursor m/z was conducted based on high-confidence (<1 % False Discovery Rate, FDR) linear peptide identifications. The re-calibrated peak lists were searched against the sequences and the reversed sequences (as decoys) of crosslinked peptides using the Xi software suite (v.1.6.745) for identification (Mendes et al., 2019). Final crosslink lists were compiled using the identified candidates filtered to 1 % FDR on link level with xiFDR v.2.1.5.2 (Lenz et al., 2021).

### Hydrogen-deuterium exchange mass spectrometry (HDX-MS)

An aliquot of 5 µl of 10 µM polymerase module or polymerase module-Mpe1 complex was incubated with 45 µl of D2O buffer at room temperature for 3, 30, 300 and 3000 seconds, and the reaction was quenched and stored at -80 °C. The quenched protein samples were rapidly thawed and subjected to proteolytic cleavage by pepsin followed by reversed phase HPLC separation. Briefly, the protein was passed through an Enzymate BEH immobilized pepsin column, 2.1 x 30 mm, 5 µm (Waters, UK) at 200 µl/min for 2 min and the peptic peptides trapped and desalted on a 2.1 x 5 mm C18 trap column (Acquity BEH C18 Van-guard pre-column, 1.7 µm, Waters, UK). Trapped peptides were subsequently eluted over 12 min using a 5-36% gradient of acetonitrile in 0.1% v/v formic acid at 40 µl/min. Peptides were separated on a reverse phase column (Acquity UPLC BEH C18 column 1.7 µm, 100 mm x 1 mm (Waters, UK). Peptides were detected on a SYNAPT G2-Si HDMS mass spectrometer (Waters, UK) acquiring over a *m/z* of 300 to 2000, with the standard electrospray ionization (ESI) source and lock mass calibration using [Glu1]-fibrino peptide B (50 fmol/µl). The mass spectrometer was operated at a source temperature of 80°C and a spray voltage of 2.6 kV. Spectra were collected in positive ion mode.

Peptide identification was performed by MS^e^ (Silva et al., 2005) using an identical gradient of increasing acetonitrile in 0.1% v/v formic acid over 12 min. The resulting MS^e^ data were analyzed using Protein Lynx Global Server software (Waters, UK) with an MS tolerance of 5 ppm.

Mass analysis of the peptide centroids was performed using DynamX sotware (Waters, UK). Only peptides with a score >6.4 were considered. The first round of analysis and identification was performed automatically by the DynamX software, however, all peptides (deuterated and non-deuterated) were manually verified at every time point for the correct charge state, presence of overlapping peptides, and correct retention time. Deuterium incorporation was not corrected for back-exchange and represents relative, rather than absolute changes in deuterium levels. Changes in H/D amide exchange in any peptide may be due to a single amide or a number of amides within that peptide. All time points in this study were prepared at the same time and individual time points were acquired on the mass spectrometer on the same day.

### *In vitro* cleavage and polyadenylation assays

To confirm the amount of enzyme used in each assay was equivalent between the complexes being compared, we analyzed 10 pmol of the complex to be used in the assay by SDS-PAGE and staining with Instant Blue (Abcam, cat. no. 119211). We measured the band intensity of Ysh1 (for cleavage assays) or Pap1 (for polyadenylation assays), normalized to the intensity of Cft1 within each respective lane using ImageJ (v. 1.52a). The ratio of normalized band intensities between complexes was used to adjust the calculated concentrations used in the assay.

Dual color cleavage assays were carried out essentially as previously described (Hill et al., 2019) with some modifications. Briefly, reactions were carried out in 1x reaction buffer (5 mM HEPES pH 8.0, 150 mM KOAc, 2 mM MgOAc, 0.05 mM EDTA) supplemented with 3 mM DTT and 1 U/μl RiboLock (Thermo). Reactions were carried out with 100 nM CF IA, 100 nM CF IB, and 50 nM of the indicated CPF complex or variant. Reactions were started by adding 100 nM of the dual-labeled *CYC1* RNA (Table S3).

Polyadenylation assays were carried out as previously described (Casanal et al., 2017; Kumar et al., 2021) under the same conditions as cleavage assays except that 2 mM ATP was added. Reactions were carried out with 100 nM 5ʹ FAM-labeled pre-cleaved *CYC1* RNA (Table S3), 50 nM enzyme as indicated, and 100 nM or 450 nM of CF IA and CF IB as indicated.

For each time point in both cleavage and polyadenylation assays, 15 μl of the reaction were stopped with 4 µl of stop solution (130 mM EDTA, 5% SDS, 12 mg/ml proteinase K in 1x reaction buffer) for 5 minutes at 37°C. Samples were mixed with 20 µl of loading buffer (1 M NaCl, 0.1 mg/ml bromophenol blue, 0.1 mg/ml xylene cyanol, 1 mM EDTA, 78% formamide). Reaction products (15 µl of prepared sample) were analyzed on a 10% (polyadenylation assays) or 20% (cleavage assays) urea-PAGE gel.

For both cleavage and polyadenylation assays, gels were scanned in a Typhoon FLA 7000 (GE Healthcare) using the 473 nm laser and Y520 filter for FAM. For cleavage assays, gels were also scanned using the 635 nm laser with the R670 filter for AlexaFluor647. Images were background subtracted (rolling ball radius of 100 pixels with light background) and subject to linear contrast enhancement in ImageJ.

For cleavage reactions, we measured the integrated intensity of the uncleaved RNA band (FAM channel) at each time point throughout the reaction. For each time point (*t*), we calculated the % of substrate cleaved relative to the initial RNA intensity (*t=0*):

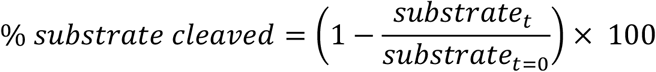

The data was fitted with a global non-linear regression using an exponential plateau function using Prism 8 (v. 8.1.2):

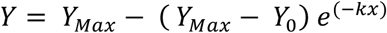

Where YMax and Y0 are globally restrained. The time it took for the enzyme to consume 50% of the maximum substrate consumed by CPF (t50) was calculated by solving for x:

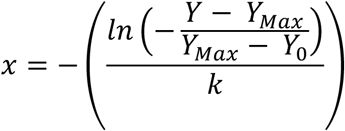

where 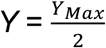,and using the following YMax, Y0 and k values for each complex:

|  | Complex |  |  |  |
| --- | --- | --- | --- | --- |
| | CPF | CPFΔMpe1 | CPF<sup>W257A/Y260A</sup> | CPF<sup>P215G</sup> |
| Best-fit values |  |  |  |  |
| $Y_{Max}$ | 83.42 | 83.42 | 83.42 | 83.42 |
| $Y_0$ | 0.8474 | 0.8474 | 0.8474 | 0.8474 |
| $k$ | 0.4412 | 0.0459 | 0.0324 | 0.0148 |
| $k$ (95% CI) | 0.3719 to 0.5291 | 0.03882 to 0.05410 | 0.02680 to 0.03882 | 0.01122 to 0.01867 |
| $R^2$ | 0.93 | 0.97 | 0.94 | 0.94 |

$Y_{Max}$ : maximum value at plateau (% of initial substrate)
$Y_0$ : starting value (% of initial substrate)
$k$ : rate constant ( $\text{min}^{-1}$ )
CI: confidence interval

### Yeast strains

Yeast strains are listed in Table S4. To construct JRY101 (Mpe1-mAID) the mAID insert was amplified from pST1933 (Tanaka et al., 2015) using Phusion high-fidelity DNA Polymerase (NEB, cat. no. M0530S), transformed into YMK728 (WT) as previously described (Chen et al., 1992), and selected on YEPD plates supplemented with 500 μg/ml G418 (Sigma, A1720-5G). Colonies were screened by PCR as previously described (Looke et al., 2011) and tested for auxin sensitivity (0.25 mM–10 mM) in growth curve assays using the Tecan 96-well plate reader with shaking at 30°C. JRY114 was generated by plating 20 µl JRY101 cells grown overnight in rich non-selective media (YEPD) onto synthetic complete media supplemented with 5-FOA (Zymo Research, cat. No. F9001-5) (Boeke et al., 1987). Colonies were verified to be URA3- and G418-resistant.

### Whole cell extract and immunoblots

Alkaline whole cell extracts were prepared as previously described (Kushnirov, 2000). 5–15 µl of whole cell extract were loaded on 4–12% Bis-Tris SDS-PAGE (Thermo, NP0323BOX) and transferred to nitrocellulose membrane (Whatman Protran BA85, cat. No. 10401196) (25 V, 1 A, 3 0min using Trans Blot Turbo transfer system (Bio-Rad). Membrane was blocked with 5% milk and probed with anti-mAID (1:5,000, MBL Life Science, cat. No. M214-3) or anti-GAPDH-HRP (1:50,000, Invitrogen, cat. No. MA5-15738-HRP). Membranes were visualized with 0.5x ECL (Amersham, cat. No. RPN2106) using the Gel Doc XR+ system (Bio-Rad).

### RNA preparation, sequencing and analysis

#### Auxin-induced depletion of Mpe1

10 ml of WT (YMK728) or Mpe1-mAID (JRY101) cells were grown overnight in YEPD + 300 µg/ml G418 at 30°C with 180 rpm shaking. Overnights were subcultured in 60 ml of YEPD to an OD600 ∼0.2 and grown to OD600 ∼0.9. Cultures were split and treated with either 1 mM auxin (Sigma, cat. No. I3750-100G-A) or an equivalent volume of 100% ethanol (solvent control) for 30 minutes. 5 mM 4-thiouracil (4tU) (Sigma, 440736-1G) was added to each condition during the final 6 minutes of auxin treatment. Cells were harvested by centrifugation at 3,000 g for 3 minutes, supernatant discarded and cell pellet flash frozen in liquid nitrogen.

#### Spike-in control

*S. pombe* cells for use as a spike-in control were prepared in advance. 10 ml of overnight wild type cells (JU60) were subcultured into 500 ml of YES media and grown to an OD600 of ∼0.8. 4tU was added to a final concentration of 5 mM and allowed to incorporate into nascent transcripts for 6 minutes. Cells were harvested and resuspended in PBS to a cell density of 15 OD/ml. 1 ml aliquots were prepared and cell pellets were flash frozen for later use.

#### RNA extraction

Cells from each strain and condition were first resuspended in 0.5 ml TRI reagent each (ThermoFisher, AM9738). One aliquot of frozen *S. pombe* was resuspended in 1 ml TRI reagent and split between the two conditions (*i.e.* 0.5 ml per -/+ auxin) of a given replicate. RNA was chloroform extracted and isopropanol precipitated following TRI reagent manufacturer instructions. Extracted RNA was DNaseI treated (NEB, M0303S) for 1 hour at 37°C followed by phenol-chloroform extraction and isopropanol precipitation. RNA concentration was measured with a Nanodrop. RNA quality was assessed with the Agilent 2100 Bioanalyzer using the RNA 6000 Nano Kit (cat. no. 5067-1511). RNA of good quality (RIN ≥ 7) was used in subsequent analyses.

#### Nascent RNA biotinylation and purification

4tU labelled RNA was biotinylated and purified as previously described (Dolken et al., 2008) with minor modifications. To pull down biotinylated RNA we used Dynabeads Streptavidin C1 beads and eluted the biotinylated RNA with freshly prepared 5% β-mercaptoethanol.

#### First strand cDNA and qPCR

First strand cDNA synthesis was performed using SuperScript III Reverse Transcriptase (Invitrogen, 18080-093) following product instructions using 1 µg of total or nascent RNA and 2.5 µM oligodT/random hexamer oligo mix. 10 µl qPCR reactions (using 1:10 diluted cDNA) were assembled using Power SYBR Green PCR master mix (Thermo, cat. no. 4367659) following product instructions and primers listed in Table S1. qPCRs were carried out in an 384-well Agilent ViiA 7 instrument. All reactions were performed in technical triplicates and Ct values averaged. To normalize to *S. pombe* spike-in controls we first averaged the Ct values of three *S. pombe* transcripts (*S. pombe* Ctk: act-1, adh-1 and gpd-3) to obtain a single spike-in value for each condition and replicate (Ctspike-in):

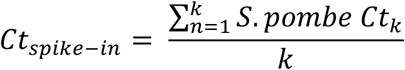

where *k* is the number of *S. pombe* transcripts (k=3).

We next normalized each Ct value from experimental *S. cerevisiae* samples to its corresponding Ctspike-in value:

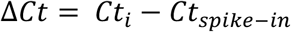

where *i* is individual replicates for each condition (*i.e.* total, nascent, -auxin, +auxin). Next we calculated the difference between +auxin and –auxin conditions, and calculated the fold change:

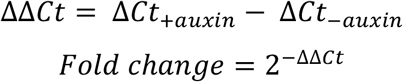

#### Library preparation and sequencing

Ribosomal RNA was depleted using the NEBNext rRNA depletion kit (NEB) following the product instructions. Depletion was confirmed with the Agilent 2100 Bioanalyzer using the RNA 6000 Nano kit.

Total and nascent RNA libraries were prepared using the NEBNext Ultra II Directional RNA Library Prep Kit for Illumina (NEB, E7760S) following the rRNA depleted RNA protocol. Libraries were pooled and single-end sequenced (50 bp) in a HiSeq 4000 instrument (Cancer Research UK).

#### Sequencing data processing and analysis

Data was trimmed with TrimGalore 0.4.5 (-q 20) and aligned to merged reference genomes of *S. cerevisiae* and *S. pombe* using STAR (v. 2.6.0a) (Dobin et al., 2013).

Aligned reads were filtered using samtools view with the following flags: -b –q 7 -F 1284. Strand-specific read quantification per genomic feature was carried with the featureCounts program of Rsubread (v. 2.0.1) (Liao et al., 2014), using the filtered reads and the merged genomic features of *S. cerevisiae* and *S. pombe*. The output of featureCounts was used with the RUVg method of RUV-seq (v. 1.20.0) (Risso et al., 2014) which used the mapped reads from *S. pombe* spike-ins to remove unwanted variability in the data. This spike-in corrected data was then used in DESeq2 (v. 1.26.0) (Love et al., 2014) (lfcThreshold = 0, alpha = 0.05, pAdjustMethod = “fdr”) to identify differentially expressed genes in the *S. cerevisiae* data.

For visualization of the data in Integrated Genome Viewer (v. 2.4.11) (Robinson et al., 2011), filtered reads from replicates were merged using samtools merge –c –r –f followed by splitting strand-specific aligned reads with samtools view –b –F 20 for the plus strand, and –b –f 16 for the minus strand. Strand-split data was then sorted (samtools sort) and indexed (samtools index –b). To visualize the data we used deeptools bamCoverage (v. 3.1.3) (Ramirez et al., 2016) to generate binned data across the genome (--normalizeUsing CPM --binSize 10 --smoothLength 20). To generate the scatter plots in Figure 6C, the output of bamCoverage was used in deeptools multiBigWigSummary bins (with --binSize 1000) or BED-file (with the ORF-T annotation (Xu et al., 2009)). Scatter plots were generated in R (v. 3.6.0). SeqPlots was used for k-means clustering (k=4) of the strand-specific nascent RNA-seq data within a 1kb window centered around the polyadenylation site. SeqPlots (Stempor and Ahringer, 2016) was also used to generate the metagene plots in Figure 6B.

**Table S1:**
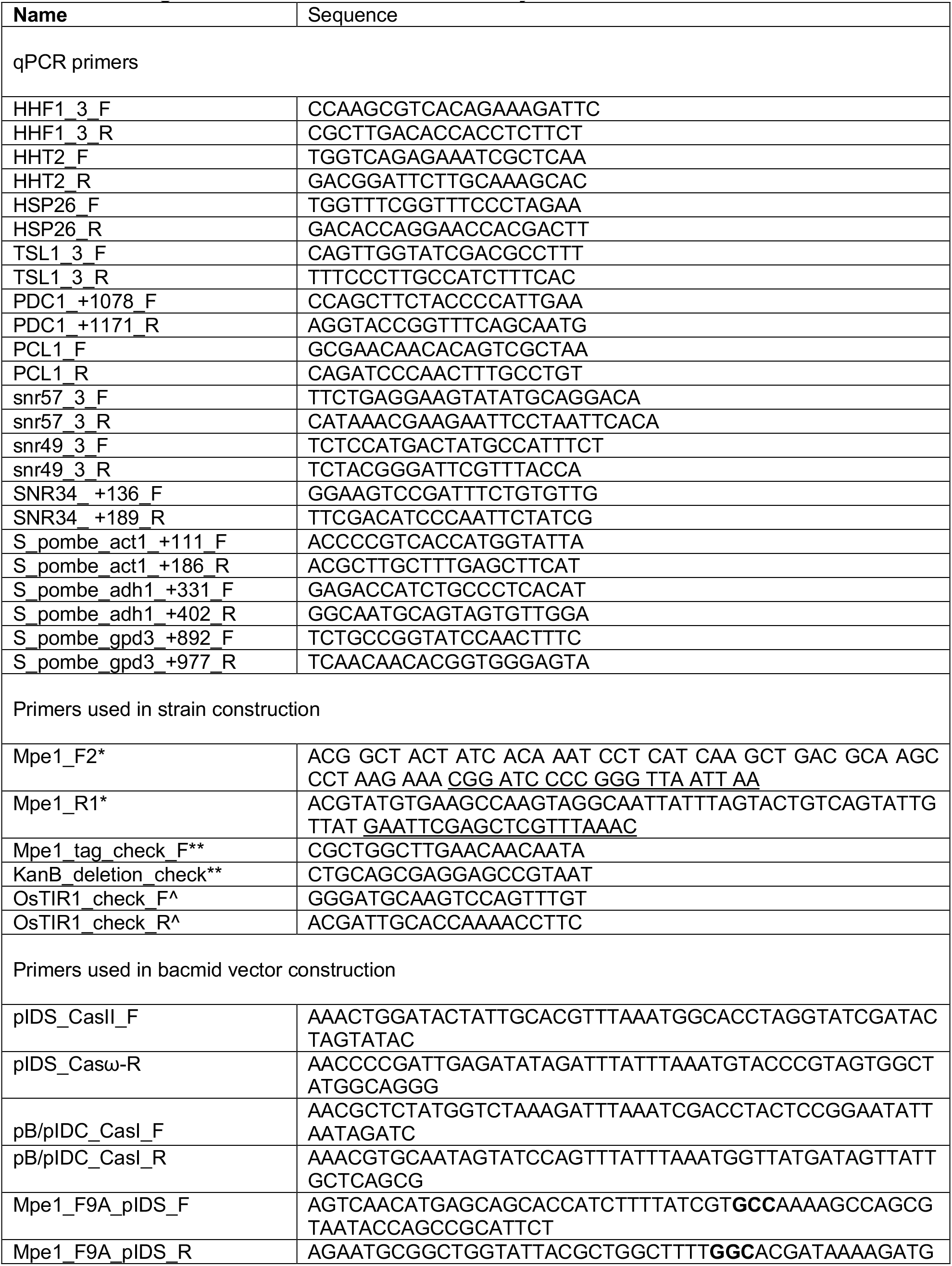
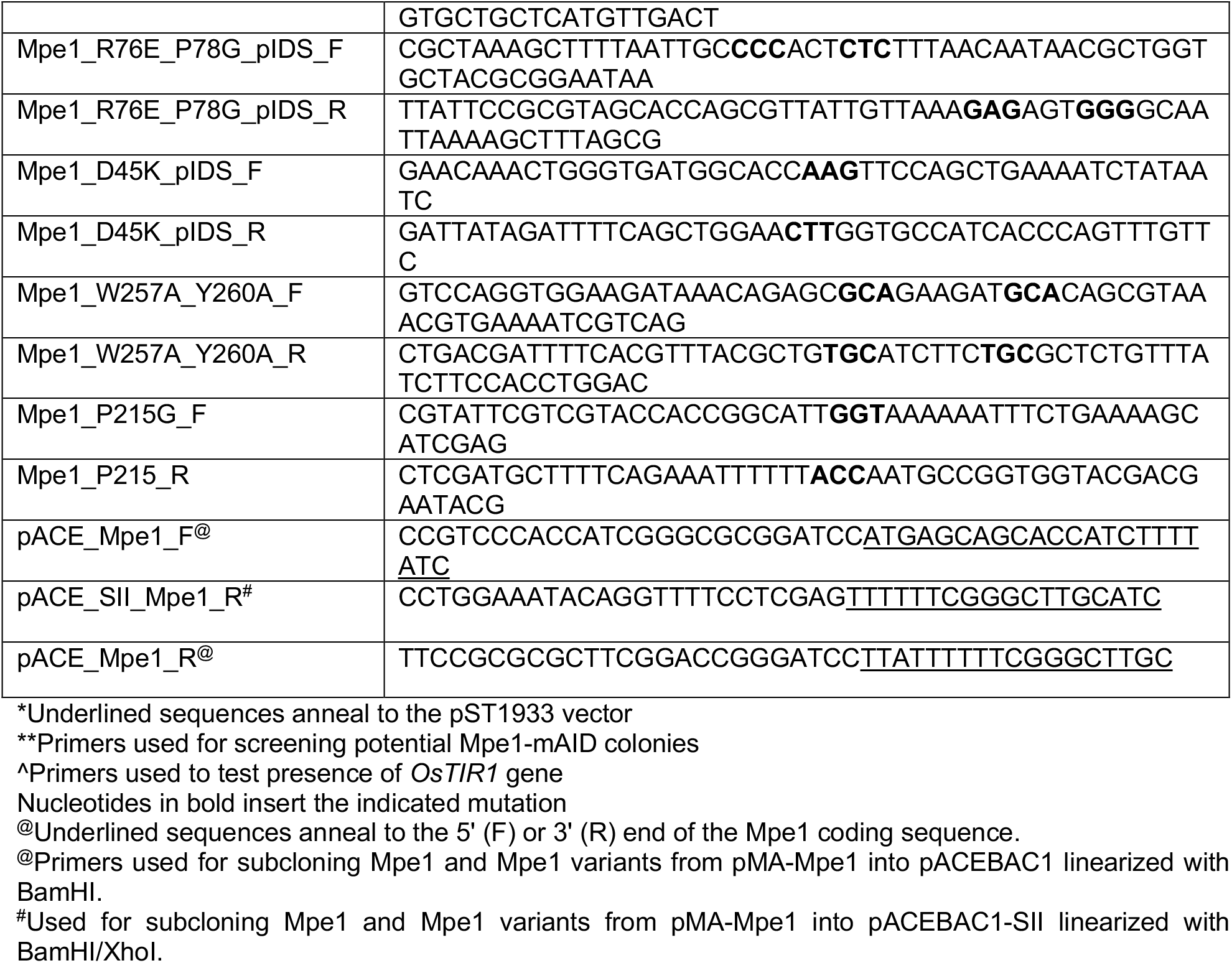
Oligonucleotides used in this study.

**Table S2:** Plasmids used in this study.

| Name | Assembled by | Source |
| --- | --- | --- |
| pBig1A-Cft1-Mpe1-SII(C) | Gibson | This study |
| pBig1A-Pap1-Mpe1-SII(C) | Gibson | This study |
| pBig1A-Pfs2-Mpe1-SII(C) | Gibson | This study |
| pBig1A-Fip1-Mpe1-SII(C) | Gibson | This study |
| pBig1A-Yth1-Mpe1-SII(C) | Gibson | This study |
| pBig1A Construct A (Cft1-Pap1-Pfs2-Fip1-Yth1) | Gibson | (Hill et al., 2019) |
| pBig1A Construct B (Cft1-Pap1-Pfs2-SII©-Fip1-Yth1) | Gibson | (Hill et al., 2019) |
| pBig2AB Construct A (Cft1-Pap1-Pfs2-Fip1-Yth1) + Mpe1-©(C) | Gibson | This study |
| pBig1B Construct AX (Ysh1-Cft2) | Gibson | This study |
| pBig2AB Ssu72-Pti1-Glc7-Re©SII(C)-Swd2 | Gibson | (Kumar et al., 2021) |
| pET28a +(modified) 6H-3C-Cft2(S) | PCR insert into NdeI/BamHI site | Chris Hill |

**Table S3:**
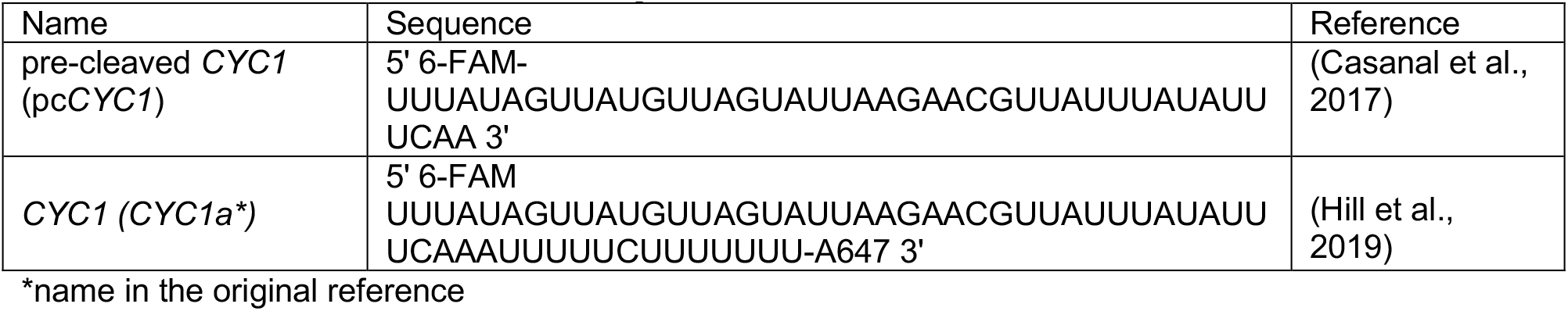
RNAs used in this study.

**Table S4:** Yeast strains used in this study.

| Strain name | Genotype | Source |
| --- | --- | --- |
| YMK728 | MATa ade2-1 his3-11,15 trp1-1 leu2-3,112 can1-100 ura3-1::ADH1-OsTIR1(pMK200, URA3) | (Nishimura and Kanemaki, 2014) |
| JRY101 | YMK728 Mpe1-3miniAID-3FLAG | This study |
| JRY114 | YMK728 Mpe1-3mAID-3FLAG (OsTIR1-, URA3-) | This study |
| JU60 | h+ | A kind gift from Dr J Mata |

---

**Figure S1.**
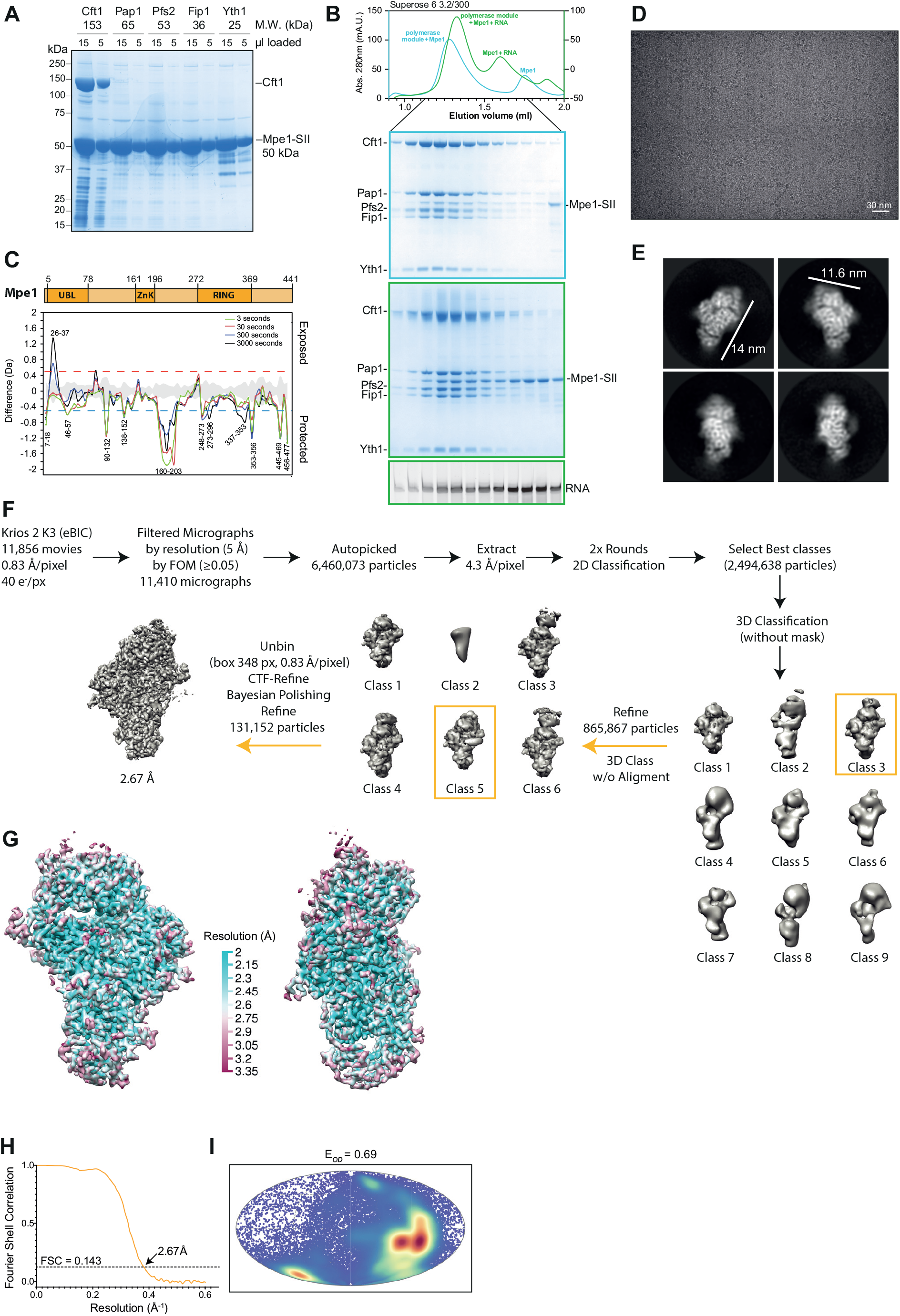
Mpe1 interacts with the polymerase module. Related to Figure 1. (A) SDS-PAGE of pulldowns from insect cell lysates co-expressing StrepII-tagged (SII) Mpe1 (bait) with individual subunits of the polymerase module. For each pulldown, two different amounts of eluate were loaded as indicated. Expected molecular weights of each subunit are indicated. (B) Co-purification of Mpe1 with the polymerase module. Size exclusion chromatography was performed for polymerase module mixed with Mpe1-SII without (blue) or with (green) a 5ʹ FAM- labelled pre-cleaved *CYC1* RNA substrate. Overlaid chromatograms are shown in the top panel. Fractions were analyzed by SDS-PAGE for protein (middle panels) and by denaturing urea-PAGE for RNA (bottom). Borders of gels correspond to colors of chromatogram traces. (C) Hydrogen-deuterium exchange mass spectrometry difference plot (Mpe1-polymerase module versus Mpe1) showing peptides of Mpe1 that are protected (negative) and exposed (positive) by interaction with the polymerase module. A domain diagram of Mpe1 is shown above. (D) Representative cryoEM micrograph of polymerase module-Mpe1-RNA complex. (E) 2D class averages of the polymerase module-Mpe1-RNA complex. (F) Schematic of data collection, processing and map reconstruction strategy for the polymerase module-Mpe1-RNA complex (see Methods for details). The nominal resolution of the final map is shown. Maps were flipped on the Z-axis following final refinement to match correct stereochemistry before model building. (G) Local resolution distribution of polymerase module-Mpe1-RNA map. (H) Fourier shell correlation (FSC) plot of sharpened cryoEM map of the polymerase module-Mpe1-RNA complex. (I) Mollweide projection of orientation distribution of particles in the final cryoEM map of polymerase module-Mpe1-RNA complex. Efficiency of orientation distribution (E*OD*) is 0.69, which indicates a minor Fourier space gap (Naydenova et al., 2017).

**Figure S2.**
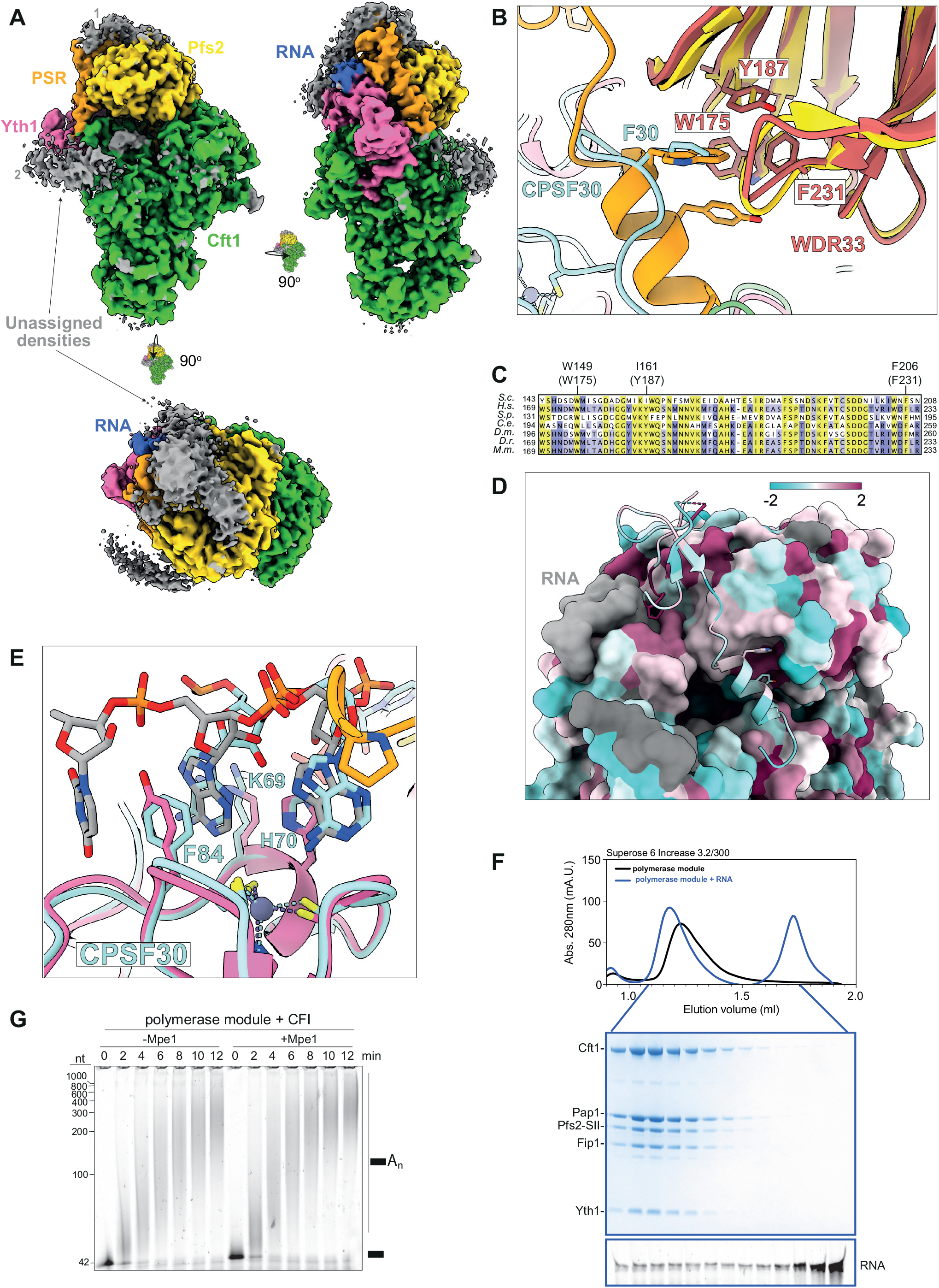
Mpe1 senses the PAS on the polymerase module. Related to Figures 1 and 2. (A) CryoEM map of polymerase module-Mpe1-RNA rendered at a lower contour level to show additional densities (grey) that could not be modeled. The density on top of Pfs2 is labeled ‘1’ and the density in front of the Cft1 C-terminal helical domain is labeled ‘2’. (B) Structure alignment of mPSF (PDB 6fuw), and polymerase module-Mpe1-RNA (this study). Yeast Mpe1 (orange) and human CPSF30 (cyan) insert into the equivalent hydrophobic pocket of Pfs2/WDR33 (yellow/ salmon) via a bulky hydrophobic residue (W257 in Mpe1, F30 in CPSF30). (C) Multiple sequence alignment of Pfs2 orthologues across model eukaryotes. Residues highlighted in yellow are conserved across most eukaryotes; those in purple are partially conserved. See Figure 1F for species abbreviations. (D) Surface rendering of the polymerase module-Mpe1-RNA model, except Mpe1 which is in cartoon representation, colored by the entropy-based conservation index implemented by AL2CO in ChimeraX. Pfs2 shows high conservation in the PSR helix binding site and near the N-terminal region of the PSR. (E) Structure alignment of RNA-bound zinc finger 2 of CPSF30 of mPSF (PDB 6fbs), and Yth1 of the polymerase module-Mpe1-RNA structure (this study). The recognition mechanism of A1A2 in the PAS is nearly identical in yeast and human. (F) Size exclusion chromatogram of polymerase module (black) or polymerase module in complex with 5ʹ FAM-labeled pre-cleaved *CYC1* RNA (blue) (top). Protein and RNA from the indicated fractions of the polymerase module-RNA complex were analyzed on SDS-PAGE and urea-PAGE, respectively (bottom panels). The polymerase module-RNA complex elutes earlier than the polymerase module alone, consistent with complex formation. (G) Polyadenylation assays similar to Figure 2E, except that 100 nM CF IA and CF IB were included in the reaction. CF IA and CF IB stimulate polyadenylation activity to comparable levels in both complexes.

**Figure S3.**
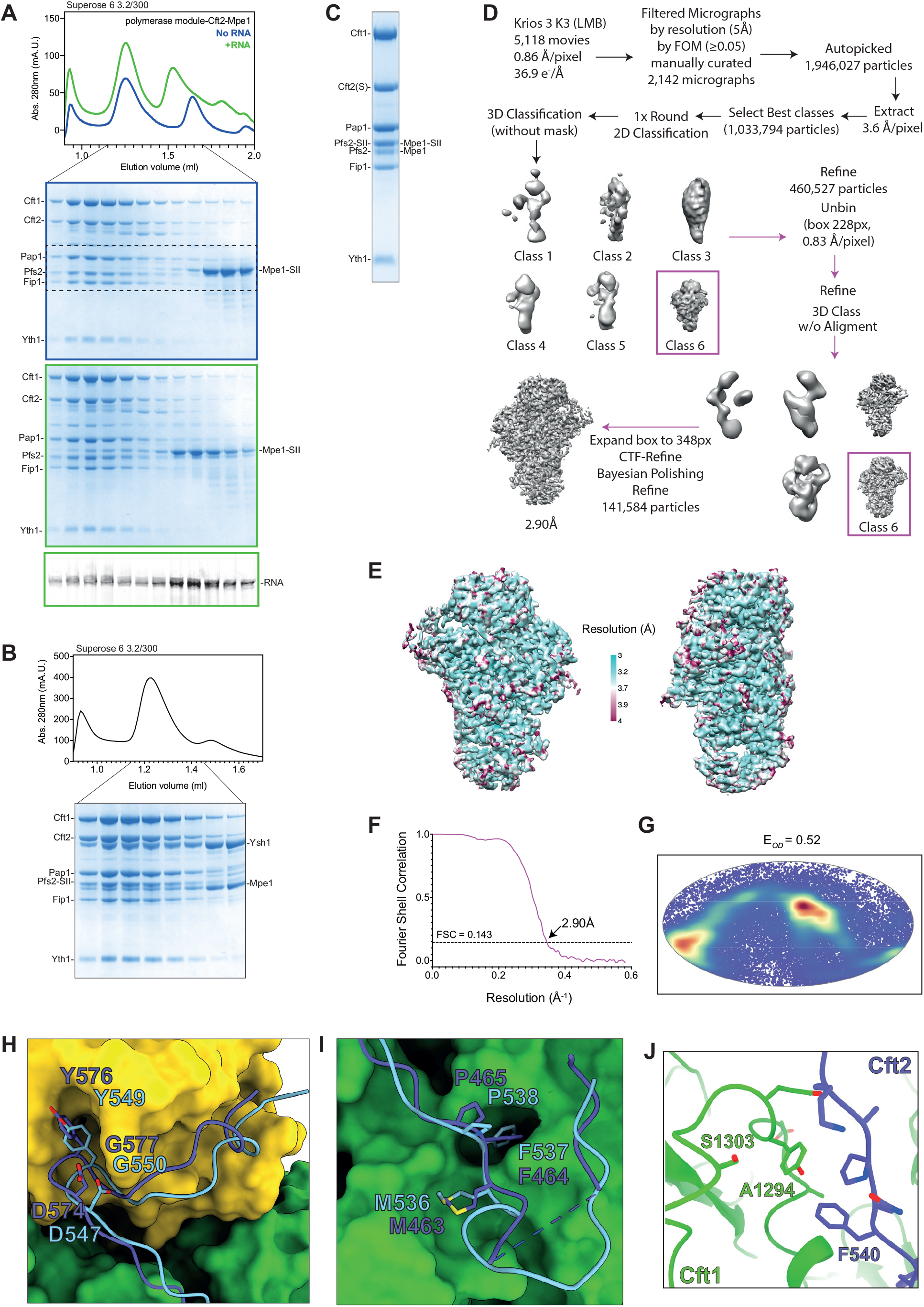
CryoEM of polymerase module-Cft2-Mpe1-RNA. Related to Figure 3. (A) Corresponding uncropped gels for Figure 3A. (B) A complex of purified Ysh1-Mpe1 was added in excess to polymerase module and analyzed by size-exclusion chromatography. Chromatogram (top) and Coomassie-stained SDS-PAGE (bottom) of indicated fractions are shown. This eight-subunit assembly is termed CPFcore (Hill et al., 2019). (C) Representative Coomassie-stained SDS-PAGE lane of the polymerase module-Mpe1- Cft2(S)-RNA complex used for cryoEM sample preparation. Complex was purified by size exclusion chromatography as described for the complex with full length Cft2. Carryover 3C protease from previous purification steps of Cft2(S) led to partial cleavage of the SII tag on Mpe1 and Pfs2. The shorter version of Cft2 was used because the C-terminal region causes problems with particle distribution in ice. (D) Schematic of data collection, processing and 3D reconstruction of polymerase module- Cft2(S)-Mpe1-RNA complex. Maps were flipped on the Z-axis following final refinement to match correct stereochemistry before model building. (E) Local resolution distribution of unsharpened cryoEM map of the polymerase module- Cft2(S)-Mpe1-RNA reconstruction. (F) Fourier shell correlation plot of sharpened cryoEM map of the polymerase module- Cft2(S)-Mpe1-RNA complex. Nominal resolution was determined at FSC = 0.143 threshold. (G) Mollweide projection of orientation distribution of particles in the final cryoEM map of the polymerase module-Cft2(S)-Mpe1-RNA complex. Efficiency of orientation distribution (E*OD* = 0.52) indicates a modest Fourier space gap (Naydenova et al., 2017). (H-I) Alignment of polymerase module-Cft2(S)-Mpe1-RNA structure with the mPSF-PIM structure (PDB 6urg). The most conserved residues of the yPIM (light blue) and PIM (purple) are shown. Cft1 (green) and Pfs2 (yellow) are shown in surface representation. (J) A disordered loop (S1303-A1294) of Cft1 (green) is stabilized upon binding to Cft2 (purple).

**Figure S4.**
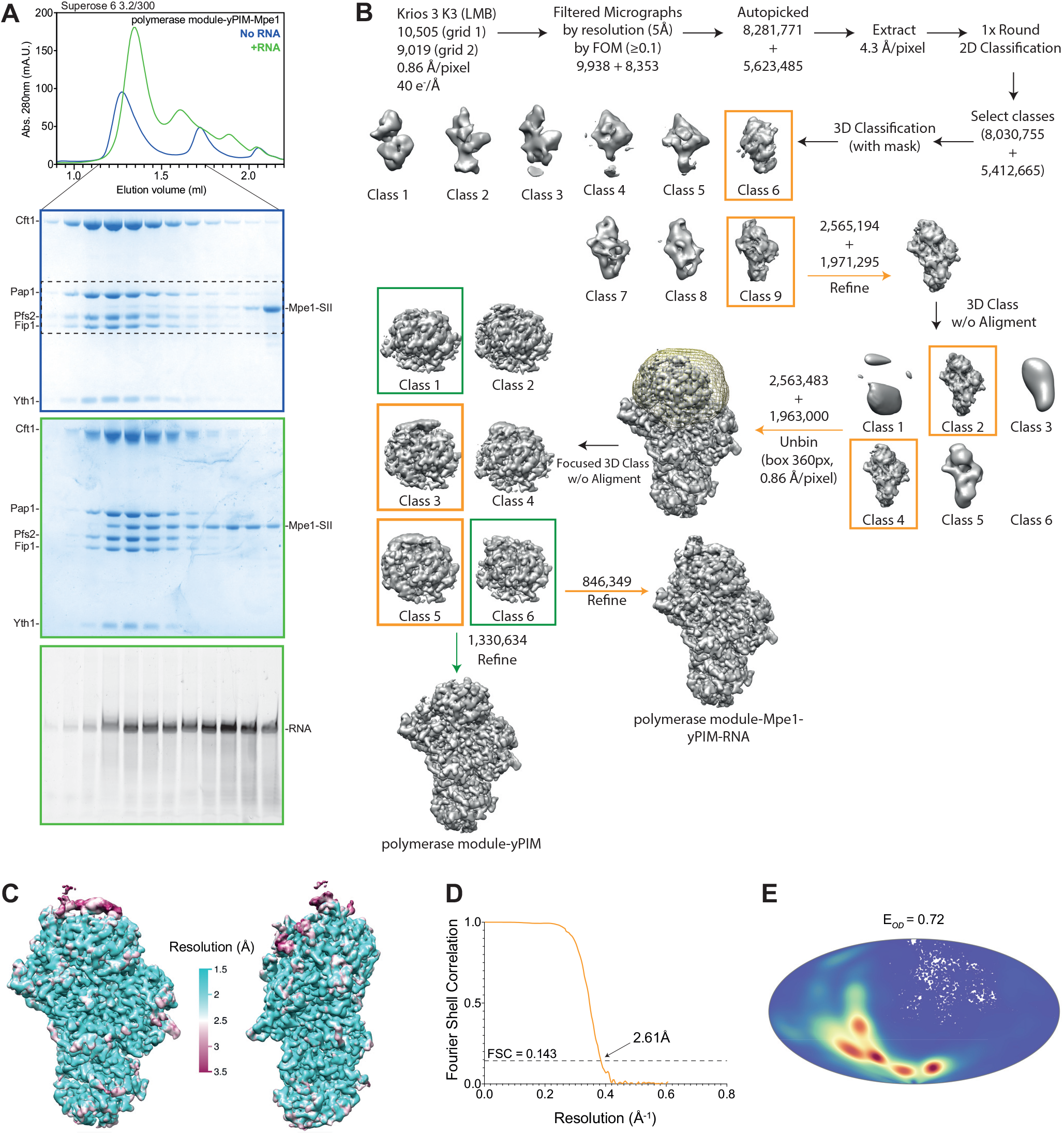
CryoEM of the polymerase module-yPIM-Mpe1-RNA complex. Related to Figure 4. (A) Size exclusion chromatography was performed with the polymerase module in complex with the yPIM of Cft2 and Mpe1 either without (blue) or with (green) pre-cleaved *CYC1* RNA. The chromatogram (top), Coomassie-stained SDS-PAGE of indicated fractions (middle two panels) and urea-PAGE gel of fluorescently labeled RNA from the indicated fractions (bottom panel) are shown. Outline color of gels correspond to the colors of the chromatogram profiles. (B) Schematic of cryoEM data collection and processing for the polymerase module-Mpe1- yPIM-RNA complex. Data was collected from two grids, and the particle number processed from each is indicated separately until merging. Focused classification with a mask around the top region of the map (yellow mesh) was necessary to parse through particles that contained or lacked clear density for the Mpe1 PSR. Selected classes are indicated with orange (with Mpe1 PSR density) or green (without Mpe1 PSR density) boxes. (C) CryoEM map of polymerase module-Mpe1-yPIM-RNA complex colored by local resolution. (D) Fourier shell correlation plot of sharpened cryoEM map of the polymerase module-Mpe1- yPIM-RNA complex. Nominal resolution was determined at FSC = 0.143 threshold. (E) Mollweide projection of orientation distribution of particles in the final polymerase module- Mpe1-yPIM-RNA cryoEM map. Efficiency of orientation distribution (E*OD* = 0.72) indicates minor Fourier space gap.

**Figure S5.**
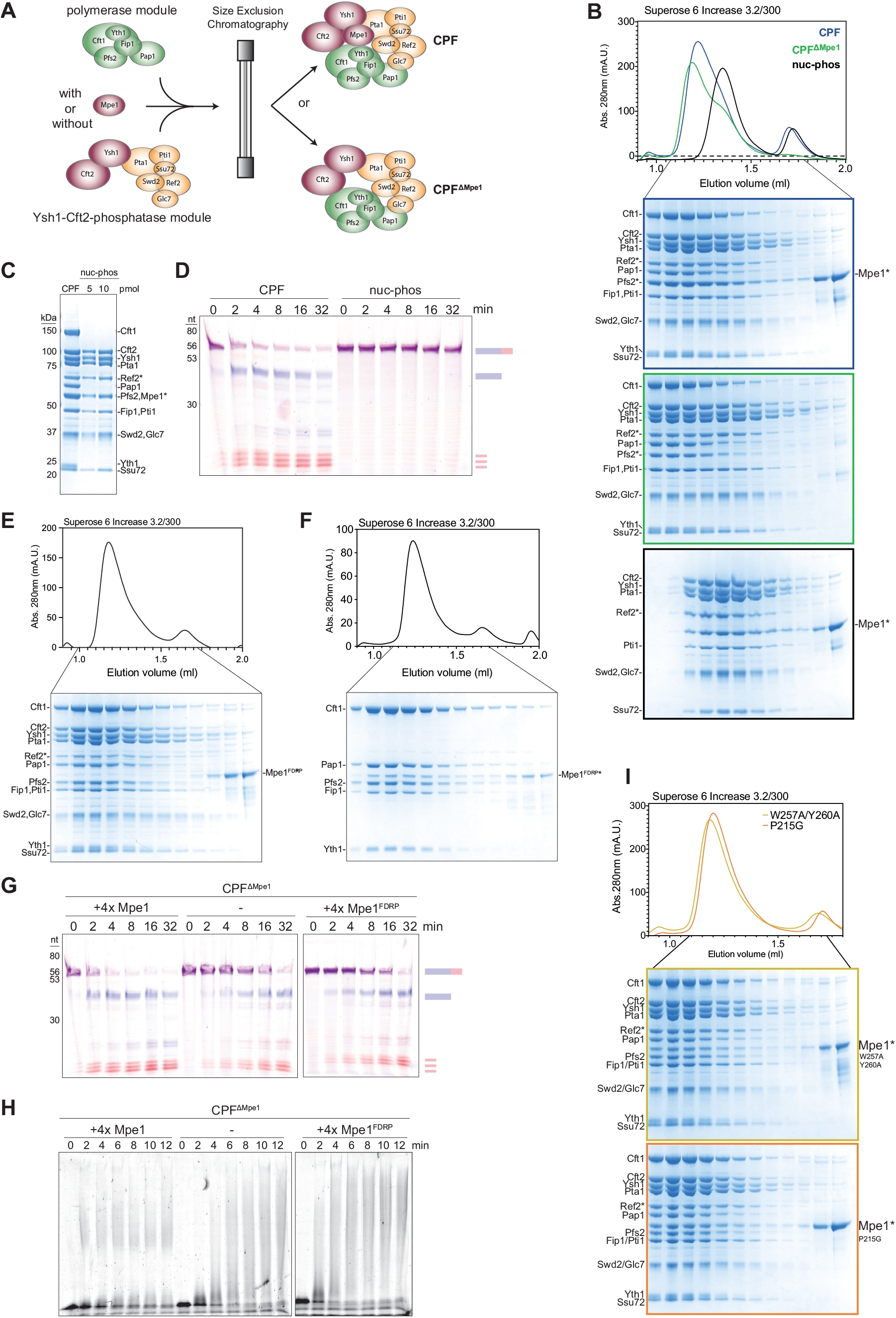
Purification and assay of recombinant CPF carrying variants of Mpe1. Related to Figure 5. (A) Schematic of CPF reconstitution and purification via size exclusion chromatography. Purified polymerase module was combined with a complex carrying all the subunits of the phosphatase module (Pta1, Pti1, Ssu72, Ref2-SII, Swd2 and Glc7) and two subunits of the nuclease module (Ctf2 and Ysh1). The assembled complex was then purified by size exclusion chromatography. The same procedure allowed us to reconstitute full CPF including Mpe1. (B) Size exclusion chromatogram of reconstituted CPF (blue), CPF^ΔMpe1^ (green) or CPF lacking the polymerase module (nuc-phos, black) is shown in the top panel. SDS-PAGE of indicated fractions for each of the reconstituted complexes are shown below. Border colors of gels correspond to chromatogram trace colors. Subunits with an asterisk (*) are SII-tagged. (C) SDS-PAGE of 10 pmol CPF, and 5 or 10 pmol of the nuc-phos module complex as indicated. Subunits with an asterisk (*) are SII-tagged. (D) Representative urea-PAGE of a dual-color *in vitro* cleavage assay using a 5ʹ FAM and 3ʹ Alexa647-labeled uncleaved *CYC1* RNA substrate with reconstituted CPF or CPF lacking the polymerase module (nuc-phos). (E-F) Size exclusion chromatogram (top) and corresponding SDS-PAGE (bottom) of CPF with an Mpe1 UBL mutant (Mpe1^FDRP^; F9A, D45K, R76E, P78G) (E) or polymerase module with Mpe1^FDRP^ (F). Mpe1^FDRP^ did not stably integrate into CPF. Subunits with an asterisk (*) are SII- tagged. (G) *In vitro* dual-color cleavage assays using CPF^ΔMpe1^. Wild-type Mpe1 or Mpe1^FDRP^ were added to the reaction in *trans* as indicated (4x the concentration of CPF^ΔMpe1^ in the reaction corresponds to 200 nM Mpe1). (H) *In vitro* polyadenylation assays using 5ʹ FAM-labeled pre-cleaved *CYC1* substrate and CPF^ΔMpe1^. WT or Mpe1^FDRP^ were added to the reaction in *trans* as indicated (4x the concentration of CPF^ΔMpe1^ in the reaction corresponds to 200 nM Mpe1). (I) Size exclusion chromatogram (top) and SDS-PAGE of indicated fractions (bottom) for CPF assemblies using purified Mpe1^W257A/Y260A^ or Mpe1^P215G^. Border colors of gels correspond to chromatogram trace colors. Both Mpe1 variants incorporate and co-elute with CPF. Subunits with an asterisk (*) are SII-tagged.

**Figure S6.**
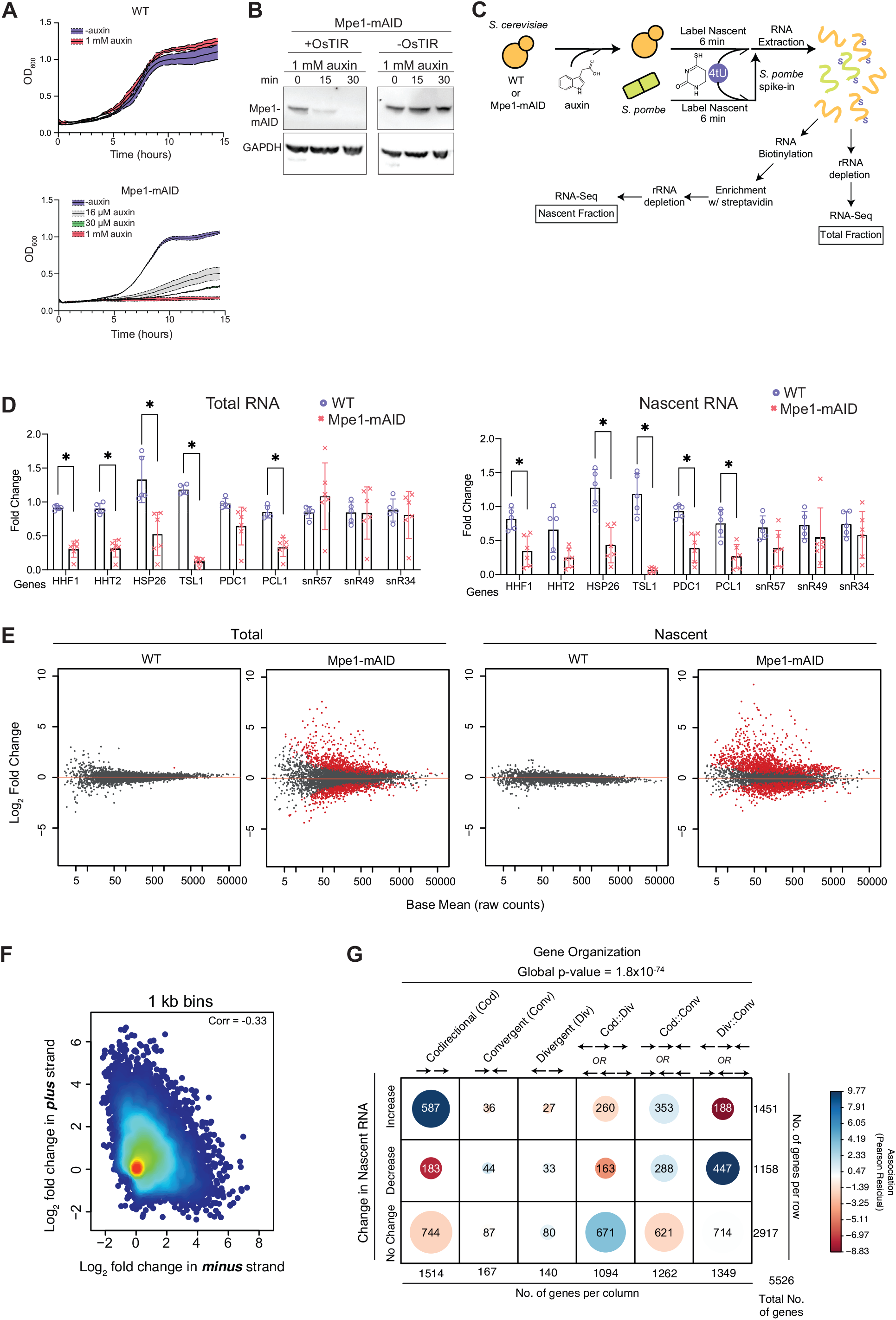
Depletion of Mpe1 results in transcription readthrough. Related to Figure 6. (A) Growth curves of WT or Mpe1-mAID cells grown in the indicated concentrations of auxin. Cultures were grown in three independent biological replicates. Standard deviations for each condition are shown as shaded areas around the mean (solid line in the middle). (B) Immunoblots of whole cell extracts from cells expressing or lacking the *OsTIR1* gene, treated with 1 mM auxin for the indicated times. Membranes were probed using antibodies against the mAID tag (top panels) or GAPDH as loading control (bottom panels). (C) Schematic of RNA labeling, and total or nascent RNA library preparation and sequencing. (D) Fold change in selected transcripts from total (left) or nascent (right) RNA fractions upon treatment of WT (purple circles) or Mpe1-mAID (red x) cells with 1 mM auxin for 30 min. N = 5 or 6 as indicated. *, P-value<0.05 Student’s T-test. (E) MA-plot of transcripts from total or nascent RNA fraction from WT or Mpe1-mAID cells as indicated. Genes that show a significant change in abundance (|log2 fold change > 0|, FDR- adjusted p-value<0.05) are highlighted in red. (F) Density scatter plot of the strand-specific log2 fold change in nascent RNA upon Mpe1 depletion. Points on the scatter plot correspond to the log2 fold change within a moving 1 kb bin across the genome on each strand. (G) *χ*^2^ test of independence between changes in nascent RNA upon Mpe1 depletion and gene organization. Genes that increase, decrease or do not change (rows) were classified based on their relationship to their neighboring gene (columns; codirectional (cod), convergent (conv), divergent (div)). Within each row, genes that are shared between two different columns (i.e. cod, conv, div) were also classified into hybrid categories (cod::div, cod::conv, div::conv). The number of genes in each classification is illustrated by the size of the circle and the number inside. The color of each circle represents the association between individual row and column categories (Pearson residual; blue or positive values denote positive association, red or negative values denote negative association).

